# Pathogen-derived 9-methyl sphingoid base is perceived by a lectin receptor kinase in Arabidopsis

**DOI:** 10.1101/2021.10.18.464766

**Authors:** H. Kato, K. Nemoto, M. Shimizu, A Abe, S. Asai, N. Ishihama, T. Daimon, M. Ojika, K. Kawakita, K. Onai, K. Shirasu, M. Ishiura, D. Takemoto, Y. Takano, R. Terauchi

## Abstract

In plants, many invading microbial pathogens are recognized by cell-surface pattern recognition receptors (PRRs), inducing defense responses; yet how PRRs perceive pathogen sphingolipids remains unclear. Here, we show that the ceramide Pi-Cer D from a plant pathogenic oomycete *Phytophthora infestans* triggers defense responses in Arabidopsis. Pi-Cer D is cleaved by an Arabidopsis apoplastic ceramidase, NCER2, and the resulting 9-methyl-branched sphingoid base is recognized by a plasma membrane lectin receptor-like kinase, RDA2. Importantly, 9-methyl-branched sphingoid base, which is unique to microbes, induces plant immune responses by interacting with RDA2. Loss of *RDA2* or *NCER2* function compromised Arabidopsis resistance against an oomycete pathogen, indicating that these are crucial for defense. We provide new insights that help elucidate the recognition mechanisms of pathogen-derived lipid molecules in plants.

**One Sentence Summary:** Oomycete-derived ceramide is cleaved into sphingoid base by ceramidase and recognized by an Arabidopsis receptor kinase.

## Main Text

Plant defend themselves against a multitude of microbial pathogens by sensing pathogen invasion through cell-surface pattern recognition receptors (PRRs) that recognize microbe- or pathogen-associated molecular patterns (MAMPs, PAMPs) or damage-associated molecular patterns (DAMPs) of host-derived molecules that emanate from damage caused by pathogen attack. This recognition then activates immune signaling *(1*-*3)*. The Arabidopsis genome contains genes encoding ∼580 PRRs, including ∼410 receptor-like kinases (RLKs) and ∼170 receptor-like proteins (RLPs) that lack the kinase domain *(4)*. However, molecular interactions between MAMP/PAMPs and PRRs have been demonstrated only in a limited number of cases, with the majority involving pathogen peptides, proteins, and carbohydrates. It has recently been reported that lipids derived from pathogens are also recognized by plant PRRs. The Arabidopsis PRR LIPOOLIGOSACCHARIDE-SPECIFIC REDUCED ELICITATION recognizes medium-chain 3-hydroxy fatty acids of bacterial pathogens *(5)*, but whether other PRRs recognize pathogen lipids remains unknown.

Ceramides belong to a class of sphingolipids consisting of a sphingoid base and a fatty acid and are present at high concentrations in eukaryotic cell membranes. Ceramide and its metabolites are also involved in intracellular signal transduction in animal cells and plants *(6*-*7)*. Recently, a ceramide-related compound, *<u>P</u>hytophthora <u>i</u>nfestans* <u>cer</u>amide <u>D</u> (Pi-Cer D; Fig. 1A), from the oomycete pathogen *P. infestans* was shown to induce immune responses in potato plants *(8)*. Pi-Cer D also induced defense responses in Arabidopsis; therefore, we aimed to identify the Arabidopsis components involved in the perception of Pi-Cer D in Arabidopsis. For this, we employed Lumi-Map, a platform consisting of a luciferase (LUC)-based mutant screen and gene identification (fig. S1) *(9)*. Because the Arabidopsis *WRKY33* gene is induced by PAMPs, including flg22, a peptide derived from bacterial flagellin and is required for resistance against pathogens *(10*-*11)*, we tested whether Pi-Cer D induced the expression of a *LUC* transgene driven by the *WRKY33* promoter (p*WRKY33*-*LUC*) in Arabidopsis, which resulted in a transient induction of bioluminescence (fig. S2). We then screened 10,000 M_2_ seedlings generated by ethylmethanesulfonate (EMS) mutagenesis of the p*WRKY33*-*LUC* reporter line (W33-1B) for mutants that showed a reduction in bioluminescence after Pi-Cer D treatment (named *L*ow (L) mutants). We isolated nine mutants insensitive to Pi-Cer D (L-09, L-12, L-16, L-19, L-31, L-46, L-55, L-66, and L-74) and two mutants with an extremely low response to Pi-Cer D (L-53 and L-107) (Fig. 1B and fig. S3, Table S1). These mutants showed normal bioluminescence responses to other PAMPs, such as flg22, elf18, derived from bacterial elongation factor Tu, and chitin, a component of fungal cell walls, indicating that they carried lesions affecting the signaling pathway that is specifically required for the response to Pi-Cer D (Fig. 1C and fig. S4). To identify the gene(s) altered in these mutants, we performed MutMap analysis *(12)*. All nine Pi-Cer D-insensitive mutants showed SNP-index peaks on chromosome 1 and contained SNPs within the gene *At1g11330* encoding a lectin receptor-like kinase RDA2 (*r*esistant to *D*FPM-inhibition of *A*BA signaling 2) (Fig. 1D, fig. S5, and Table S2), a mutant of which (*rda2*) is insensitive to a small synthetic molecule [5-(3,4-dichlorophenyl)furan-2-yl]-piperidine-1-ylmethanethione (DFPM) and incapable to mount DFPM-mediated immune signaling and inhibition of ABA signaling *(13)*. Thus, we tentatively named the nine Pi-Cer D-insensitive mutants as *rda2-4* through *rda2-10*. The two Pi-Cer D low-response mutants showed SNP-index peaks on chromosome 2 and carried mutations in the gene *At2g38010*, which encodes neutral ceramidase 2 (*NCER2*, Fig. 1E, fig. S5, and Table S2) *(14)*. We then tentatively named L-53 and L-107 mutants as *ncer2-2* and *ncer2-3*, respectively. Complementation of the *rda2* and *ncer2* mutant lines by the respective wild-type alleles restored bioluminescence induction following Pi-Cer D treatment, confirming that *RDA2* and *NCER2* are the responsible genes for the given phenotypes (Fig. 1F and fig. S6). Furthermore, T-DNA insertion mutant lines for *RDA2* and *NCER2* showed either no or reduced induction of *WRKY33* gene expression after Pi-Cer D treatment (fig. S7). Collectively, these results indicate that *RDA2* and *NCER2* are required for Pi-Cer D recognition in Arabidopsis. We then asked whether *RDA2* and *NCER2* contribute to Arabidopsis immunity against an oomycete pathogen *Hyaloperonospora arabidopsidis*. Importantly, both *rda2* and *ncer2* mutants showed increased susceptibility to *H. arabidopsidis* (Fig. 1G), indicating that *RDA2* and *NCER2* are required for resistance to this pathogen.

**Fig. 1.**
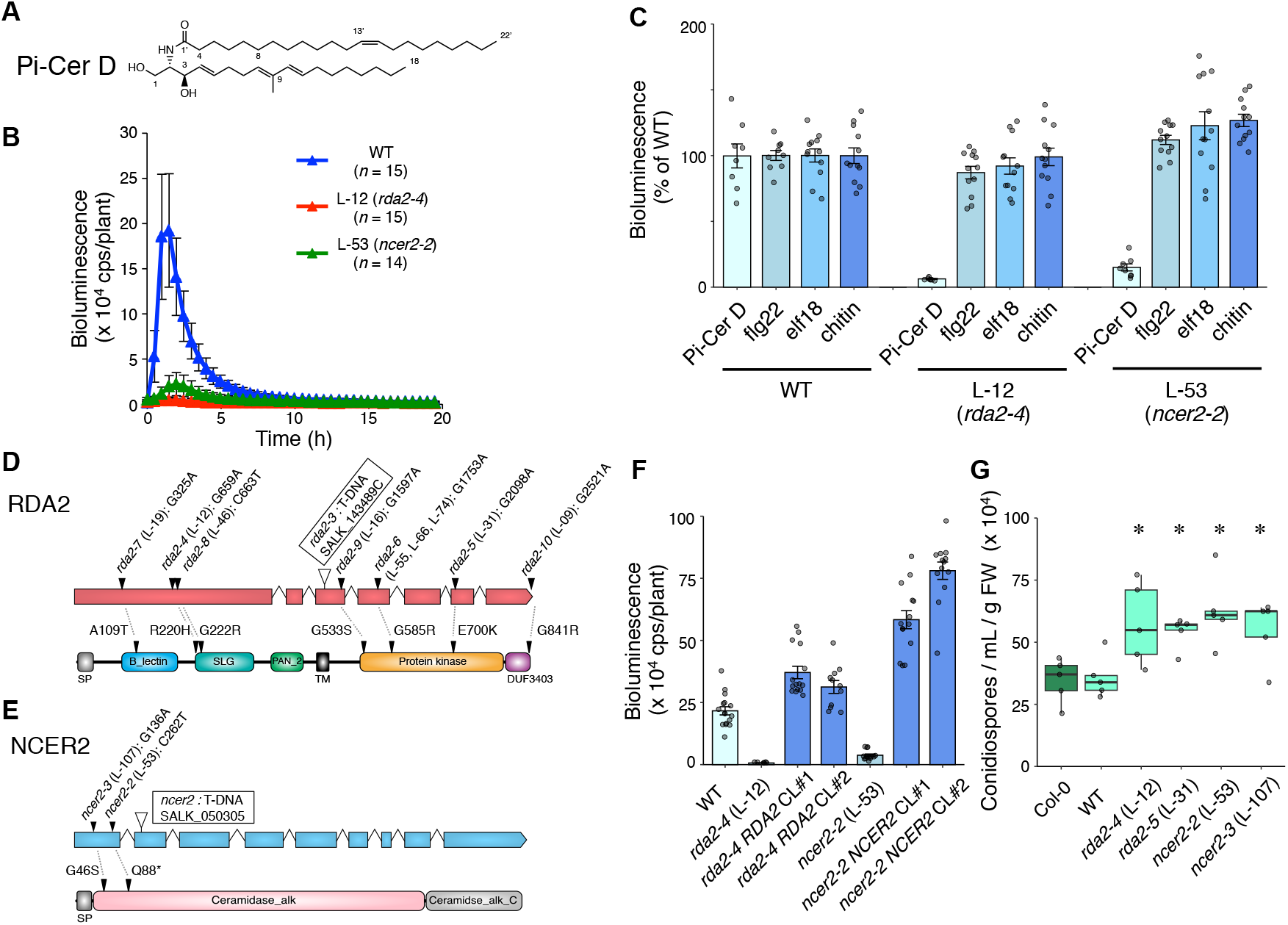
Arabidopsis *RDA2* and *NCER2* are required for recognition of Pi-Cer D and resistance against *Hyaloperonospora arabidopsidis*. (**A**) Chemical structure of Pi-Cer D. (**B**) Bioluminescence response over time of Arabidopsis p*WRKY33*-*LUC* reporter (WT), *rda2-4*, and *ncer2-2* after Pi-Cer D (0.17 μM) treatment (means ± SD). For additional data, see fig. S3. (**C**) Bioluminescence of WT and mutant seedlings after treatment with Pi-Cer D, flg22, elf18, or chitin. Relative peak bioluminescence values are shown as % of WT (means ± SE). For additional data, see fig. S4. (**D** and **E**) Gene and protein structures of RDA2 (**D**) and NCER2 (**E**). Gene structure (top), showing exons in boxes and introns as lines between the boxes. Protein structure (bottom), showing the different domains. The positions of the EMS-induced point mutations in different alleles (closed triangle) and T-DNA insertion sites (opened triangle) are indicated. (**F**) Complementation of *rda2* and *ncer2* mutants with wild-type alleles. Bioluminescence (means ± SE) of WT, *rda2-4, ncer2-2*, and complemented lines (CL) after Pi-Cer D (0.17 μM) treatment is shown. For additional data, see fig. S6. (**G**) Growth of *Hyaloperonospora arabidopsidis* on Arabidopsis Col-0, WT, and *rda2* and *ncer2* mutants. Three-week-old Arabidopsis plants were inoculated with *Hpa* Waco9. Conidiospores were harvested and counted 5 days post inoculation (*n* = 5). *, *p* < 0.05 in two-tailed *t*-tests comparing the corresponding values from Col-0. Experiments were performed three times with similar results.

We hypothesized that (i) Pi-Cer D is cleaved by NCER2 into a mature ligand product in the apoplastic space and (ii) the ligand is recognized via plasma-membrane-localized RDA2. To test the first hypothesis, we investigated whether the *ncer2* mutant phenotype was rescued by the product generated by NCER2 ceramidase treatment of Pi-Cer D (Fig. 2A). The lipid fraction containing Pi-Cer D and NCER2 produced in *Nicotiana benthamiana*, as well as that containing Pi-Cer D and a mouse ceramidase, induced p*WRKY33*-*LUC* bioluminescence in the *ncer2-2* mutant (L-53) (Fig. 2B). We observed no bioluminescence when we used NCER2^G46S^, a mutant version of NCER2 present in *ncer2-3* (L-107). These results indicate that Pi-Cer D was cleaved by *NCER2*-encoded ceramidase and the resulting compound was recognized by RDA2. The predicted molecular size of NCER2 tagged with hemagglutinin (HA-NCER2) was 82 kDa; however, the protein detected by immunoblot analysis using anti-HA antibody was 26 kDa (fig. S8). Upon purifying the HA-NCER2 protein and subjected it to gel electrophoresis, we recovered two protein bands (26 kDa and 56 kDa) (fig. S8). Analysis of the bands by liquid chromatography-tandem mass spectrometry (LC-MS/MS) revealed that they corresponded to the N- and C-terminal regions of NCER2 (Table S3), respectively. These results indicate that NCER2 is processed into its N- and C-terminal regions, which function together. To investigate the localization of NCER2, we then generated transgenic Arabidopsis *ncer2-2* mutant lines that expressed HA-NCER2 driven by its own promoter (*ncer2-2 HA-NCER2*) (fig. S9). We detected the HA-NCER2 protein in the apoplast wash fluid (AWF) of these lines (Fig. 2D) and found that *WRKY33-LUC* activity was induced in the AWF from the wild-type reporter line and *ncer2-2 HA-NCER2* lines, but not in that from the *ncer2-2* mutant (Fig. 2C). These results indicate that NCER2 localized to the apoplast and metabolizes Pi-Cer D into a mature ligand product that is recognized by RDA2.

**Fig. 2.**
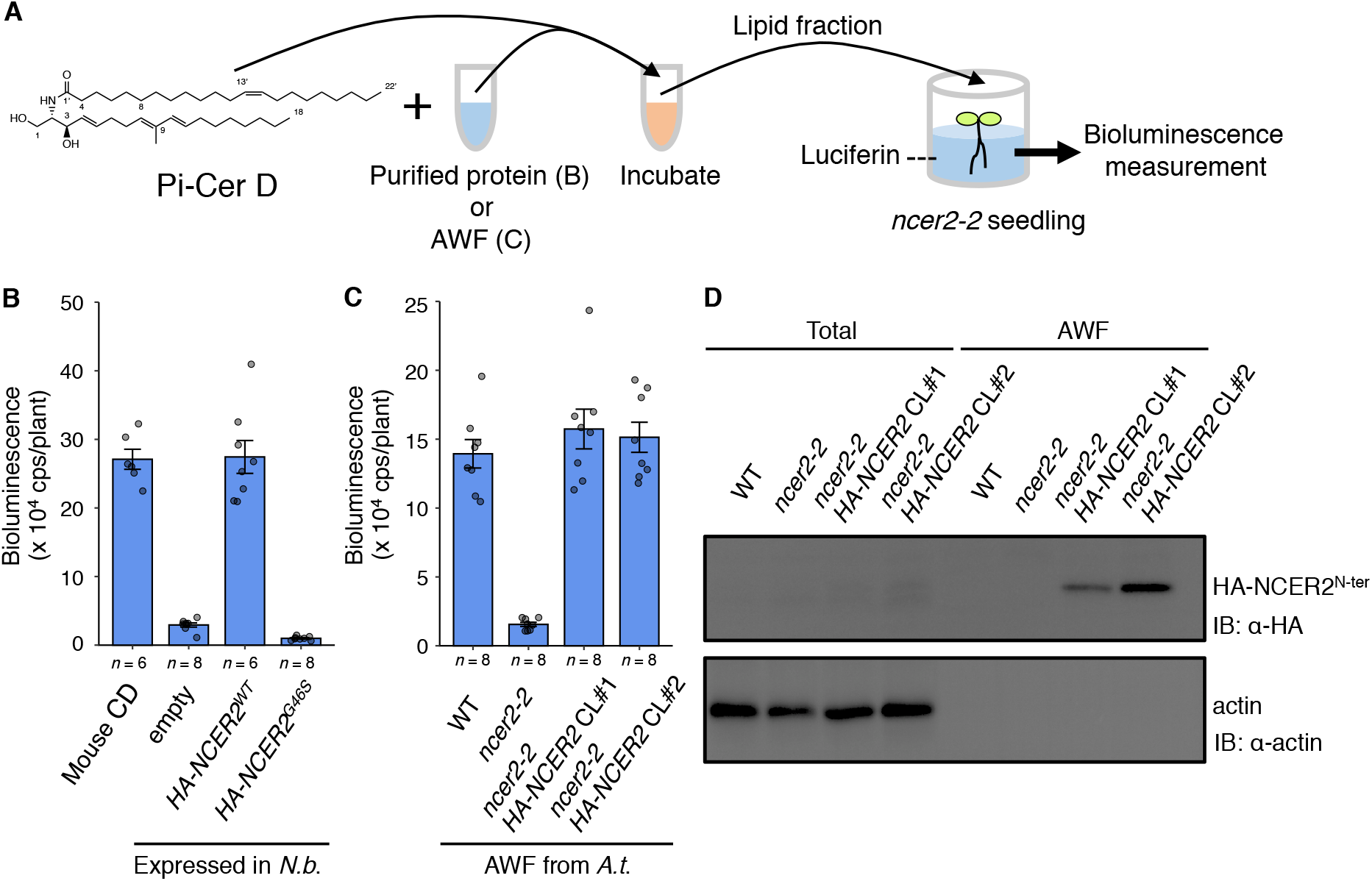
Pi-Cer D is cleaved by NCER2, an apoplastic ceramidase. (**A**) Pi-Cer D-cleavage assay. Pi-Cer D was incubated for 24 h with HA-NCER2, its mutant variant produced in *Nicotiana benthamiana* leaves, or apoplast wash fluid (AWF) from Arabidopsis plants. The lipid fraction containing metabolites derived from Pi-Cer D was recovered and applied to *ncer2-2* plants, and their bioluminescence was measured. (**B**) Pi-Cer D is cleaved by the Arabidopsis ceramidase NCER2. HA-tagged wild-type (HA-NCER2^WT^) and mutated NCER2 (HA-NCER2^G46S^, carrying the same mutation as in the L-107 line) were transiently expressed in *N. benthamiana* (*N*.*b*.) and purified. Commercial mouse ceramidase (Mouse CD) served as a positive control. Peak bioluminescence is shown (means ± SE). For additional data for NCER2 expressed in *N. benthamiana*, see fig. S8. (**C**) Pi-Cer D-cleavage activity of Arabidopsis (*A*.*t*.) AWF. AWFs were isolated from the p*WRKY33*-*LUC* reporter line (WT), *ncer2-2* and the *ncer2-2 HA-NCER2* complementation lines (CL). Peak bioluminescence is shown (means ± SE). For additional data with *ncer2-2 HA-NCER2* plants, see fig. S9. (**D**) Immunoblot analysis of total protein and AWF extracted from WT, *ncer2-2* and *ncer2-2* HA-NCER2 plants. Anti-actin antibody was used to detect cytosolic protein. Experiments were performed three times with similar results.

Mass spectrometry analysis of compounds in the lipid fraction prepared from a mixture of Pi-Cer D and ceramidase detected a sphingoid base, suggesting that the sphingoid base derived from Pi-Cer D might be the ligand for RDA2 (fig. S10). We also compared the *WRKY33-LUC*-inducing activity of Pi-Cer D analogs, which revealed that structural differences in the sphingoid base determine the level of induction (fig. S11). The sphingoid base in Pi-Cer D ((4*E*,8*E*,10*E*)-9-methyl-4,8,10-sphingatrienine, 9Me,4*E*,8*E*,10*E*-d19:3) contains a unique branching methyl group at the ninth carbon position. Remarkably, this 9-methyl-branching structure is present in sphingoid bases of oomycete, fungi and marine invertebrates, but has not been reported in plants and mammals *(15-17)*. We thus hypothesized that the 9-methyl-branching structure of the sphingoid base is decisive in distinguishing between ‘self’ and ‘nonself’ in plants. Therefore, we investigated the ability of various sphingoid bases to induce a defense response. Because the sphingoid base in Pi-Cer D (9Me,4*E*,8*E*,10*E*-d19:3) was difficult to obtain, we used (4*E*,8*E*)-9-methyl-4,8-sphingadienine (9Me,4*E*,8*E*-d19:2, hereafter 9Me-Spd) to evaluate 9-methyl structure (Fig. 3A). Notably, the *rda2* mutants were almost insensitive to 9Me-Spd, indicating that 9Me-Spd is specifically recognized by RDA2 (fig. S12). Among the sphingoid bases we tested, 9Me-Spd showed the strongest RDA2-dependent elicitor activity (Fig. 3B, 3C, fig. S12 and S13). In addition, (4*E*,8*E*)- 4,8-sphingadienine (4*E*,8*E*-d18:2, Spd) and sphingosine (4*E*-d18:1, Sph), neither of which contain 9-methyl branching, also showed elicitor activity, although this was significantly weaker than that of 9Me-Spd (Fig. 3C, figs. S12 and S13). To identify the structural correlates of RDA2-dependent sensing of the sphingoid base, we tested sphingosine derivatives with different lengths of long-chain bases. Among these derivatives, Sph (4*E*-d18:1) induced the highest bioluminescence in the p*WRKY33*-*LUC* reporter line, followed by 4*E*-d16:1, 4*E*-d14:1, and 4*E*-d12:1 (fig. S14). This indicates that efficient sensing by RDA2 requires a long-chain base structure that includes 18 carbon atoms. We also tested phytosphingosine (4-hydroxysphinganine, 4-t18:0, PHS) and found that it did not elicit bioluminescence in the p*WRKY33*-*LUC* reporter line. This indicates that the 4*E* double-bond structure in Sph is crucial for its sensing by RDA2 (fig. S15).

**Fig. 3.**
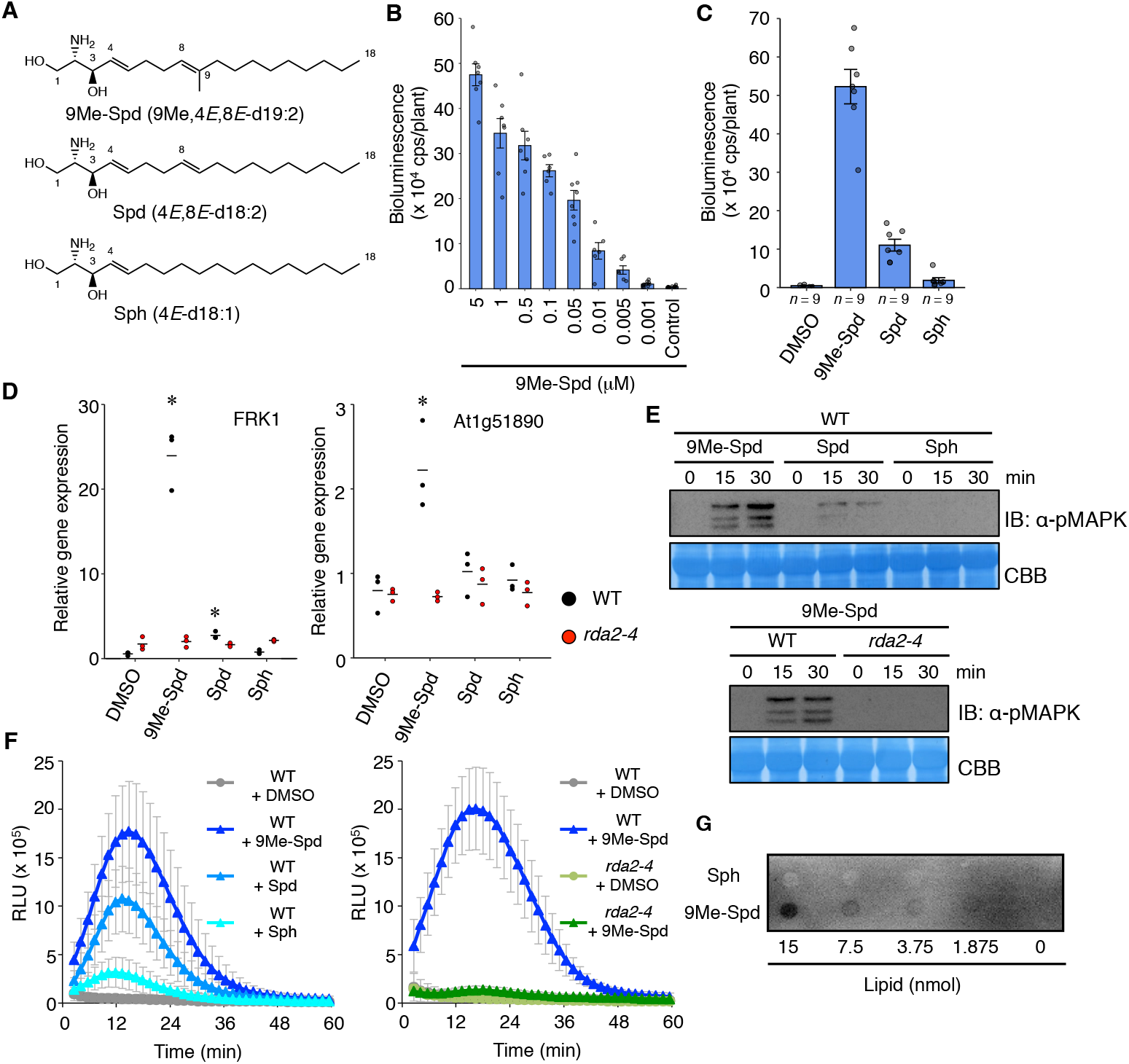
RDA2 recognizes 9-methyl sphingoid base and induces defense responses. (**A**) Structures of (4*E*,8*E*)-9-methyl-4,8-sphingadienine (9Me-Spd), (4*E*,8*E*)-4,8-sphingadienine (Spd), and sphingosine (Sph). (**B** and **C**) Peak bioluminescence values for Arabidopsis p*WRKY33-LUC* reporter seedlings (WT) treated with (**B**) different concentrations of 9Me-Spd or (**C**) 0.5 μM of structurally different sphingoid bases (means ± SE). For additional data, see fig. S12 and S13. (**D**) Expression of defense genes (*FRK1* and *At1g51890*) in Arabidopsis seedlings 3 h after elicitation with sphingoid bases (0.5 μM) relative to expression in DMSO-treated WT. Individual data (symbols) and means (bars) are shown (*n* = 3); *, *p* < 0.05 (two-tailed *t*-test). (**E**) Mitogen-activated protein kinase (MAPK) activation following treatment with sphingoid base (0.5 μM). Phosphorylated MAPKs were visualized with anti-phospho-p44/p42 MAPK antibody. (**F**) ROS accumulation in Arabidopsis leaves treated with sphingoid bases. Left, leaf disks from WT plants treated with 30 μM 9Me-Spd, Spd, or Sph (*n* = 12). Right, leaf disks from WT and *rda2-4* plants treated with 30 μM 9Me-Spd (*n* = 10). Relative light unit (RLU) is shown (means ± SD). (**G**) Binding of sphingoid bases with HA-tagged RDA2 by protein-lipid overlay assay. For additional data, see fig. S16 and S17. Experiments were performed two (**B, D**, and **F**) or three times (**C, E**, and **G**) with similar results.

To further investigate the downstream events following RDA2-mediated sensing, we tested the ability of 9Me-Spd and its derivatives to activate Arabidopsis immune responses. The 9Me-Spd activated RDA2-dependent bioluminescence induction, transcript accumulation of defense-related genes (*FRK1, At1g51890*), phosphorylation of mitogen-activated protein kinases, and the production of reactive oxygen species (ROS) more strongly than Spd and Sph (Fig. 3C–F and fig. S12). A protein-lipid overlay assay using a membrane fraction containing HA-tagged RDA2 demonstrated physical interaction between 9Me-Spd and RDA2 (Fig. 3G, fig. S16 and S17). Collectively, these results suggest that RDA2 is the receptor for sphingoid bases including 9-methyl sphingoid base, which is derived from Pi-Cer D.

Sphingolipids are major components of eukaryote membranes *(16-18)*. Our findings revealed that oomycete-derived ceramide is cleaved by plant apoplastic ceramidase and the generated sphingoid base is recognized by a lectin receptor-like kinase (Fig. 4). This indicates that plants perceive differences in sphingolipid structure for non-self recognition. Notably, plant RDA2 senses the 9-methyl-branching structure of sphingoid bases that are prevalent in oomycetes and fungi. It has recently been reported that *RDA2* is required for immune signaling and inhibition of ABA signaling by a small synthetic molecule DFPM as identified by a chemical genetic screen *(13)*. We hypothesize that DFPM or its metabolized product functions as a mimic of sphingoid base, but further study is required to clarify this. Based on our results, we propose the name *SphingR* (for *<u>sphing</u>*oid *<u>r</u>*ecognizing) as a synonym of *RDA2* (Fig. 4). Our study here provides a basis on which to engineer RDA2/SphingR to detect various pathogen-specific lipids and to enable plants mount defense against pathogens such as *P. infestans*, the causal agent of the potato late blight that devastated potato crop and caused famine in the nineteenth century.

**Fig. 4.**
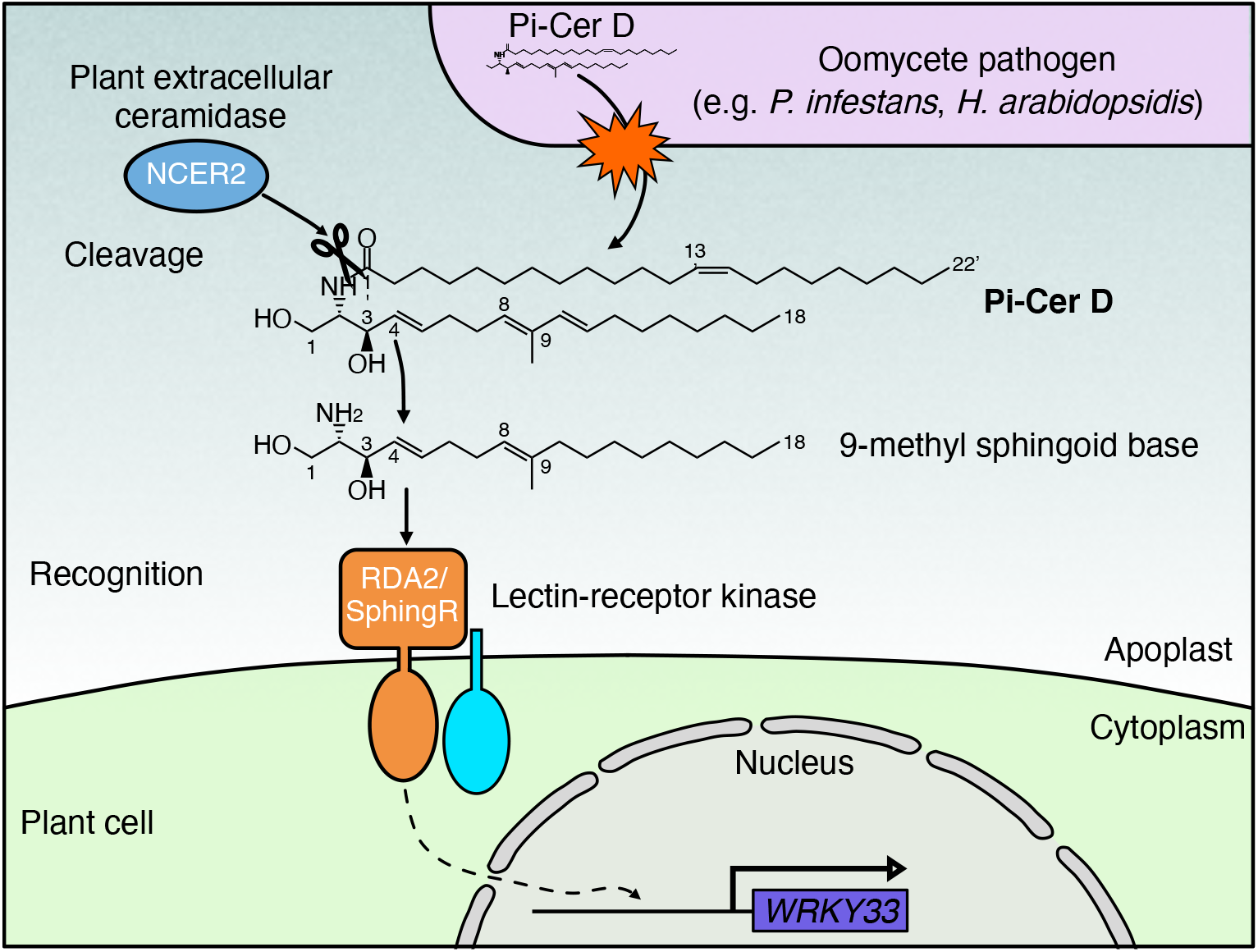
A model for the recognition of pathogen-derived ceramide in plants. Pi-Cer D is cleaved by plant apoplastic ceramidase NCER2 into 9-methyl sphingoid base. 9-methyl sphingoid base is recognized by a lectin-receptor kinase, RDA2/SphingR, which then induces defense responses that include *WRKY33* gene expression and enhances immunity against pathogen infection.

## Supporting information

Kato_Suppl_Files

## Acknowledgments

We thank H. Utsushi, E. Kanzaki, K. Ito, E. Sato (in IBRC) and Y. Inoue (in Kyoto university) for technical support. Computational analysis was partially performed on the NIG supercomputer at the ROIS National Institute of Genetics.

## Funding

This work was supported by the Japan Society for the Promotion of Science KAKENHI grants (20K15528, H.K.; 15H05779 and 20H00421, R.T.; 20H02995, S.A.; 17H06172, K.S.; 21K19112 and 21H05032, Y.T.; 17H03963, K.K; 20H02985, D.T.).

## Author contributions

H.K., D.T., Y.T., K.S., and R.T. conceived this study. H.K. performed main experiments and data analyses. M.S. and A.A. performed MutMap analysis. D.T., K.K., and M.O. purified and provided Pi-Cer D and performed HPLC analysis. K.O. and M.I. developed the bioluminescence monitoring system. S.A. and K.S. performed inoculation assays. K.N, N.I, T.D., and K.S. performed protein expression and binding assays. H.K. drafted the manuscript. H.K., D.T., Y.T. K.S., and R.T. wrote the manuscript.

## Competing interests

The authors declare no conflicts of interest in relation to this work.

## Data and materials availability

All data are available in the main text or supplementary materials.

## Supplementary Materials

Materials and Methods

Figures S1 – S17

Tables S1 – S6 References (*19* – *26*)

## Notes

### Competing Interest Statement

The authors have declared no competing interest.

