## Supplementary material for "Pathogen-derived 9-methyl sphingoid base is perceived by a lectin receptor kinase in Arabidopsis": Kato_Suppl_Files

#### Materials and Methods

Plant materials and growth conditions

All *Arabidopsis thaliana* lines are in the Col-0 ecotype background. Seeds of *rda2-3* (SALK_143489C) and *ncer2* (SALK_050305) were obtained from the Arabidopsis Biological Resource Center. Surface-sterilized seeds of Arabidopsis were incubated at 4°C in the dark for 2 days and grown in Murashige and Skoog (MS) liquid medium (1/2 MS, 0.5% sucrose, B5 vitamin solution, 2 mM MES, pH 5.7) under controlled conditions (22°C, 24-h photoperiod, 57 μmol $m^{-2}sec^{-1}$). Soil-grown Arabidopsis plant were grown on soil under controlled conditions (22°C, 10-h photoperiod, 57 μmol $m^{-2}sec^{-1}$).

Soil-grown *Nicotiana benthamiana* plants were grown individually in a pot under controlled conditions (25°C, 16-h photoperiod, 57 μmol $m^{-2}sec^{-1}$).

Molecular cloning

We confirmed the nucleotide sequences of all constructs using standard DNA manipulation and sequencing techniques. Primers used in this study are listed in Table S4.

The p*WRKY33-LUC* reporter gene cassette consisted of the *Arabidopsis WRKY33* promoter (p*WRKY33*; -2,000 to -1), the coding region of a modified firefly *luciferase* gene ($LUC^{+}$, Promega), and $T_{NOS}$. The p*WRKY33-LUC* reporter gene cassette was inserted into pBIB-HYG *(19)* with *Sal*I and *Kpn*I (pBIB-HYG-pWRKY33-LUC).

For complementation of *rda2* mutants (*rda2-4*, *rda2-5* and *rda2-6*), a 5.2 kb genomic fragment containing the *RDA2* gene and its native promoter was amplified and inserted into pBI101 with *Sal*I and *Sac*I (pBI101-RDA2) by In-Fusion HD Cloning Kit (Takara). For *ncer2* mutants (*ncer2-2* and *ncer2-3*), a 5.3 kb fragment containing *NCER2* were amplified and inserted into pBI101 with *Sal*I and *Sac*I (pBI101-NCER2). For stable transgenic plant expressing HA-tagged NCER2, *NCER2* promoter (p*NCER2*; -2,315 to -1) and *HA-NCER2* cDNA fragments were amplified and inserted into pBIB-KAN with *Sal*I and *Sac*I site (pBIB-KAN-HA-NCER2). For stable transgenic plant expressing HA-tagged RDA2, *RDA2* promoter (p*RDA2*; -2,184 to -1), *RDA2* and 3x*HA* cDNA fragments were amplified and inserted into pBIB-KAN with *Sal*I and *Sac*I site (pBIB-KAN-RDA2-3xHA).

The cloning site with 3xHA fragment from pRI 35S-HA *(20)* was inserted into *Age*I/*Xho*I site of pEAQ-HT *(21)* (pEAQ-HT-3xHA). For transient assay in *N. benthamiana*, *HA-NCER2* and *HA-NCER2*$^{G46S}$ fragments were amplified from pBIB-KAN-HA-NCER2 and inserted into pEAQ-HT-3xHA with *Age*I and *Sal*I site, respectively (pEAQ-HA-NCER2, pEAQ-HA-NCER2$^{G46S}$).

Plant transformation

Plasmids were electroporated into *Agrobacterium tumefacians* GV3101::pMP90 strains. Arabidopsis plants were transformed with the constructs by floral dipping. For constructs in pBIB-HYG, hygromycin B resistant ($Hyg^{R}$) $T_1$ plants were selected and grown using standard techniques. We selected $T_2$ plants that showed a 3:1 segregation ratio for hygromycin B resistance, suggesting they contained a single T-DNA, and obtained $T_3$ plants from these selected plants. These $T_3$ plants were selected as homozygous by lack of segregation for hygromycin B resistance. Bulk $T_4$ seeds were generated from the selected $T_3$ plants. For transformation with pBI101- or pBIB-KAN-derived vectors, kanamycin was used for selection.

Measurement of bioluminescence response

The bioluminescence response of seedlings was performed as follows. Surface sterilized seeds were incubated at 4°C in the dark for 2 days and sown into wells of a 96-well microplate (Luminunc$^{TM}$ Plates White F96; Thermo Fisher Scientific) containing 150 μL MS liquid medium containing 50 μM D-luciferin-K (Biosynth) and germinated in continuous light. After 7 days, seedlings were treated with elicitors and 96-well plates were sealed with a plate seal (Excel Scientific) instead of plastic cover.

Bioluminescence from each well was measured automatically using a commercially available automated bioluminescence monitoring system (model CL96-4; Churitsu Electric Corp.) with a robotic plate conveyor (model CI-08L; Churitsu Electric Corp.). Bioluminescence data was analyzed using commercially available software (SL00-01; Churitsu Electric Corp.).

Elicitors and Chemicals

Pi-Cer D is purified from *Phytophthora infestans* as previously described *(8)*. Other elicitors and chemicals were ordered or purchased: flg22 and elf18 peptides (Life Technologies Japan), chitin (C9752, Sigma), (4*E*,8*E*)-9-methyl-4,8-sphingadienine (9Me,4*E*,8*E*-d19:2, 9Me-Spd), (4*E*,8*E*)-4,8-sphingadienine (4*E*,8*E*-d18:2, Spd), Sphingosine (4*E*-d18:1, Sph) and Sphingosine derivatives (4*E*-d16:1, 4*E*-d14:1, 4*E*-d12:1) (Nagara Science), phytosphingosine (4-t18:0, PHS) (Tokyo Chemical Industry).

Mutagenesis of p*WRKY33-LUC* reporter line

About 5,000 $T_4$ seeds from the p*WRKY33-LUC* reporter line (W33-1B) were mutagenized by treatment with 0.15% (v/v) ethylmethanesulfonate (Merck KGaA) for 15 h at 25°C. $M_2$ seeds were collected and grouped into 24 pools, each of which contained seeds from about 200 $M_1$ plants. About 800 seedlings from each $M_2$ pool were screened for an altered bioluminescence response following Pi-Cer D treatment.

Generation of $F_2$ progeny and whole-genome sequencing

To generate the $F_2$ progeny used for bulk sequencing, each mutant was crossed to the p*WRKY33-LUC* reporter line (W33-1B, the parental line of the mutants) and the resulting $F_1$ progeny were self-pollinated to produce $F_2$ seeds. Bulk DNA for MutMap analysis was prepared from equal amounts of 30 $F_2$ mutant individuals. For whole-genome sequencing, DNA samples were extracted from young leaves with the DNeasy Plant Mini Kit (QIAGEN). We prepared 12 sequence libraries for WGS; (1), the p*WRKY33-LUC* reporter line W33-1B (read length; 150 bp, paired-end sequencing (PE); (2) L-09, L-12, L-16, L-19, L-31, L-46, L-53, L-55, L-66, L-74, and L-107 mutant lines (150 bp, PE). Sequence libraries for PE short reads were constructed using an Illumina TruSeq DNA LT Sample Prep Kit (Illumina). The libraries were sequenced on a HiSeq high output (Table S5). Whole genome sequencing data has been deposited with DDBJ BioProject under DRA012770.

MutMap analysis

Bulked segregant analysis, as implemented in MutMap *(12)* was performed using MutMap pipleline (https://github.com/YuSugihara/MutMap). As Arabidpsis reference sequence, we used the Col-0 reference sequence download from the following site (ftp://ftp.ensemblgenomes.org/pub/release-36/plants/fasta/ Arabidopsis _thaliana/dna/ Arabidopsis_thaliana.TAIR10.dna.toplevel.fa). Whole genome sequencing of the 11 independent mutants indicated that each line had 769 ± 225 (mean ± s.d.; range 479-1,113) SNPs relative to the W33-1B, the wild type reporter strain (Table S6).

RNA extraction and qRT-PCR

Gene expression analysis was performed on eight-day-old seedlings. Seedlings were treated with Pi-Cer D (0.34 μM), 9Me-Spd (0.5 μM), Spd (0.5 μM) or Sph (0.5 μM) and frozen in liquid nitrogen at indicated time. Total RNA was extracted using RNeasy Plant Mini kit (QIAGEN) according to the manufacturer's instructions. cDNA was synthesized using Takara Prime Script RT Master Mix (Takara). Takara TB Green Premix Ex Taq I and a Thermal Cycler Dice real Time System TP850 (Takara) were used for quantitative RT-PCR using the primers listed in Table S4. Relative gene expression were calculated with ΔΔCt method as described previously *(22)* and normalized against *UBC* gene.

Pathogen inoculation

*Hyaloperonospora arabidopsidis* (*Hpa*) Waco9 inoculation was done as described in Asai et al. (2015) *(23)*. Briefly, Arabidopsis plants were spray-inoculated to saturation with a spore suspension of 1 x $10^4$ conidiospores/ml. Plants were covered with a transparent lid to maintain high humidity (90-100%) conditions in a growth cabinet at 16°C under a 10-h photoperiod until the day for sampling. To evaluate conidiospore production, 5 pools of 3 plants for each Arabidopsis line were harvested in 1 ml of water at 5 dpi. After vortexing, the amount of conidiospores released was determined using a haemocytometer.

Transient expression of HA-NCER2 in *N. benthamiana* leaf

Plasmids were electroporated into *A. tumefacians* GV3101::pMP90 strains. The bacterial suspension ($OD_{600}$=0.3, 10 mM $MgCl_2$) was infiltrated into leaves of 5-week old *N. benthamiana* with a needleless syringe. After 2 days, leaves were harvested, frozen in liquid nitrogen and homogenized. Protein were extracted in extraction buffer containing 20 mM Tris-HCl, pH7.5, 50 mM NaCl, 0.25% NP-40 and 1x cOmplete [EDTA-free, Roche Applied Science]). HA-tagged NCER2 were detected by immunoblot analysis using anti-HA antibody (3F10, 1: 2000 dilution, Roche). HA-tagged protein was purified by commercial kit (HA-tagged Protein Purification kit, code 3320, MEDICAL & BIOLOGICAL LABORATORIES) according to the manufacturer's instructions.

For identification of protein bands, eluate containing purified protein was separated by SDS-PAGE and identified with silver staining (SilverQuest™ silver staining kit, Thermo Fisher Scientific). Protein bands were excised from the gel, digested with trypsin and subjected to nano-LC/MS/MS analysis by standard protocol (Japan Proteomics Co.Ltd.).

Pi-Cer D-cleaving assay

The reaction mixture contained 1 μg Pi-Cer D and 2 μl of eluate containing HA-tagged protein in 100 μl of 25 mM MES-KOH, pH 5.8, 0.25%(w/v) Triton X-100, 2.5 mM $CaCl_2$. Following incubation at 37°C for 24 h, reaction was stopped by adding 200 μL of methanol:chloroform (1:1) and the lipid fraction was dried under a SpeedVac. Equal volume of lipid was treated to *ncer2-2* mutant to detect reaction product by bioluminescence monitoring. A commercial mouse ASAH2 ceramidase (Cosmo Bio) was used as positive control of this assay.

The ceramidase reaction product derived from 10 μg Pi-Cer D was dissolved in MeOH (100 μL) and a portion (10 μL) was used for HPLC analysis under the following conditions:

column Develosil ODS-UG-5 (4.6 x 250 mm, Nomura Chemical Co., Seto, Aichi, Japan), solvent 60-100-100% (0-20-60 min) A in B (A: MeOH-2 M $NH_4OAc$ (99:1), B: 20 mM $NH_4OAc$), flow rate 1.0 mL/min, detection 245 nm. A new peak (sphingoid base) and Pi-Cer D were detected at 24.3 and 55.5 min, respectively. To collect a pure sample of the sphingoid base, HPLC was performed under the same conditions except for the solvent [90-100% (0-10 min) A in B]. The peak at 8 min was collected and analyzed by electrospray ionization mass spectrometry (ESI MS) in the positive ion mode (fig. S10).

Extraction of apoplast wash fluid

Extraction of apoplast wash fluid (AWF) was performed according to the method described by Gentzel et al. (2019) with minor modifications *(24)*. Extraction buffer (20 mM Tris-HCl, pH7.5, 50 mM NaCl) was syringe-infiltrated into leaves. After being blotted dry with a paper towel, leaves were wrapped around the 1-mL pipette tip with a parafilm and placed in 15-mL conical tube. The tube was centrifuged at 1,000 x g for 10 min in swing rotor. AWF was collected at the bottom of the tube. Protein content of AWF was measured and 500 ng AWF protein were subjected to Pi-Cer D-cleavage assay. Immunoblot analysis was performed with anti-HA (1: 2000 dilution, Roche) and anti-actin (AC009, 1: 2000 dilution, ABclonal Biotechenology) antibodies. .

MAPK activation assay

MAPK activation assays were performed on eight-day-old seedlings grown in liquid medium. Seedlings were then elicited with 9Me-Spd (0.5 μM), Spd (0.5 μM) or Sph (0.5 μM) for 15 or 30 min and frozen in liquid nitrogen. Proteins were extracted in extraction buffer (50 mM HEPES-KOH pH 7.4, 5 mM EDTA, 0.5 mM EGTA, 50 mM beta-glycerophosphate, 10 mM NaF, 10 mM $Na_3VO_4$, 2 mM DTT). MAPK activation was monitored by western blot with antibodies that recognize the dual phosphorylation of the activation loop of MAPK (pTEpY). Phospho-p44/42 MAPK (Erk1/2; Thr-202/Tyr-204) rabbit monoclonal antibodies were used according to the manufacturer's protocol (#9101, Cell Signaling Technology). Blots were stained with CBB (PageBlue™ Protein Staining Solution, Thermo Fisher Scientific) to verify equal loading.

ROS assay

ROS released by leaf tissue was assayed using 4 mm leaf discs prepared from 5-week old soil-grown Arabidopsis plant in 96 well plates. The leaf disks were pre-incubated in liquid MGRL medium *(25)* containing 1% sucrose for overnight and treated with 30 μM sphingoid base (9Me-Spd, Spd or Sph), 100 μM L-012 (FUJIFILM Wako Chemicals) and 10 μg/mL horseradish peroxidase (Sigma-Aldrich Japan K.K). Luminescence was measured as relative light unit (RLU) for 60 minutes using bioluminescence monitoring system (model CL96-4; Churitsu Electric Corp.).

Protein-Lipid overlay assay

Protein-lipid overlay assay was performed as described with minor modification *(26)*. The membrane fraction protein was isolated from Arabidopsis leaf expressing RDA2-3xHA using Minute™ Plant Plasma Membrane Protein Isolation Kit (Invent Biotechnologies) according to the manufacturer's instructions. The pellet containing the membrane fraction protein was suspended in a transmembrane protein extraction reagent (FIVEphoton Biochemicals) containing a phosphatase inhibitor cocktail (PhosSTOP, Roche) and a protease inhibitor cocktaill (Sigma-Aldrich), and the protein concentration was measured using the DC Protein Assay (Bio-Rad). The PVDF membrane (Immobilon-P, Merck) spotted with lipid was incubated with blocking buffer

(PBS-T containing 3% bovine serum albumin) containing 40 μg of membrane fraction protein at 4°C for overnight with gently shaking. To detect RDA2-3xHA, the membrane was incubated with an anti-HA-HRP (clone 3F10) antibody (1: 10000 dilution, Roche) for 1 h at room temperature with shaking. Antibody-bound proteins were detected using Immobilon ECL Ultra Western HRP Substrate (GE Healthcare) according to the manufacturer's instructions with a LAS 4000 system (GE Healthcare).

Accession numbers

The accession numbers for *Arabidopsis* genes in this article are: *RDA2/SphingR* (At1g11330), *NCER2* (At2g38010), *WRKY33* (At2g38470).

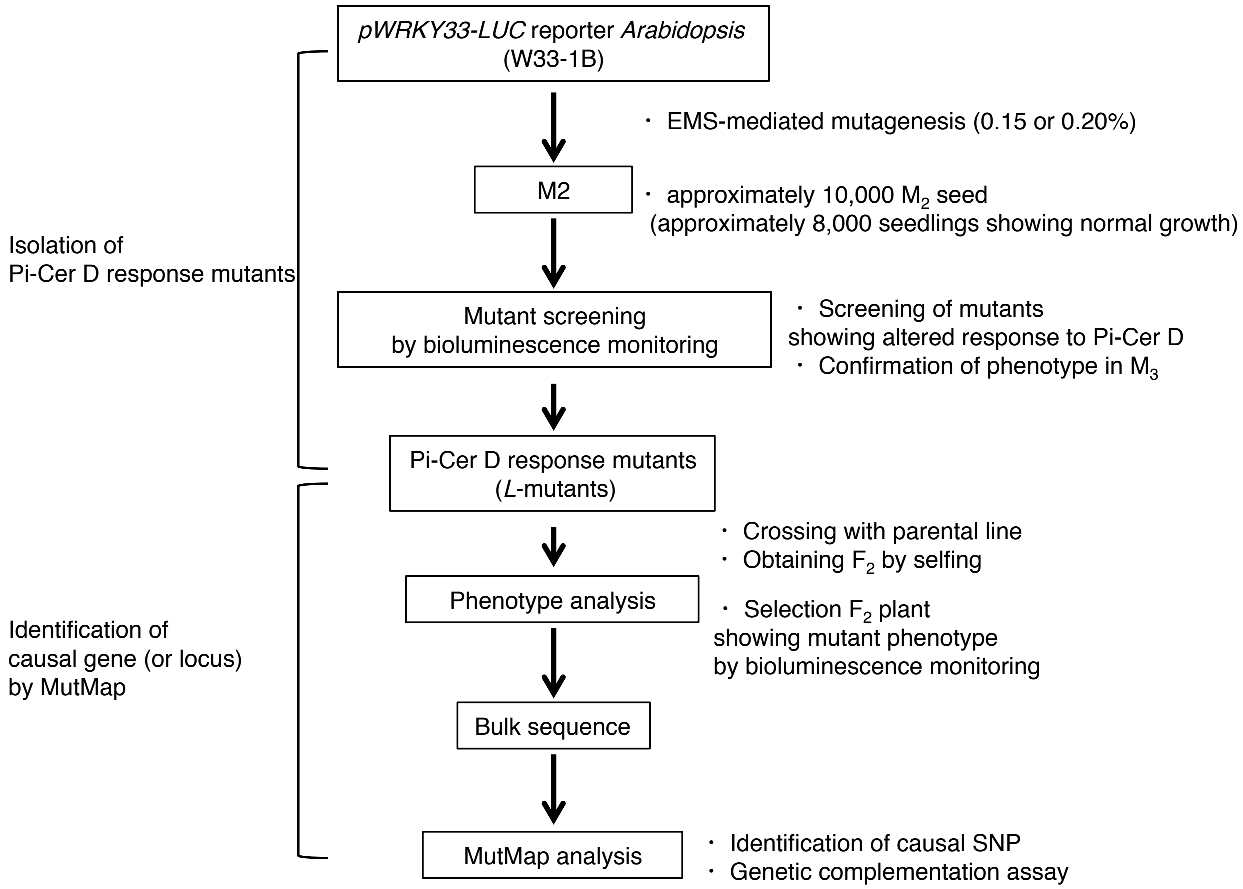


**Fig. S1. Overview of the Lumi-Map method.** A p*WRKY33-LUC* reporter strain (W33-1B) was mutagenized with ethylmethanesulfonate (EMS) and $M_2$ progeny were obtained. Mutant screening was performed by bioluminescence monitoring of Pi-Cer D-treated $M_2$ seedlings. $M_2$ seedlings that showed a mutant bioluminescence phenotype were propagated to $M_3$ and *Low* (*L*) mutants were then selected after confirmation of the bioluminescence phenotypes. Identification of the causal gene was performed using MutMap. *L* mutants were crossed with the parental reporter line (W33-1B) and $F_1$ progeny were selfed to obtain the $F_2$ progeny. $F_2$ plants were treated with Pi-Cer D and their bioluminescence phenotypes were monitored. The DNA of $F_2$ plants showing mutant phenotypes were subjected to whole-genome sequencing followed by MutMap analysis to identify causal SNPs.

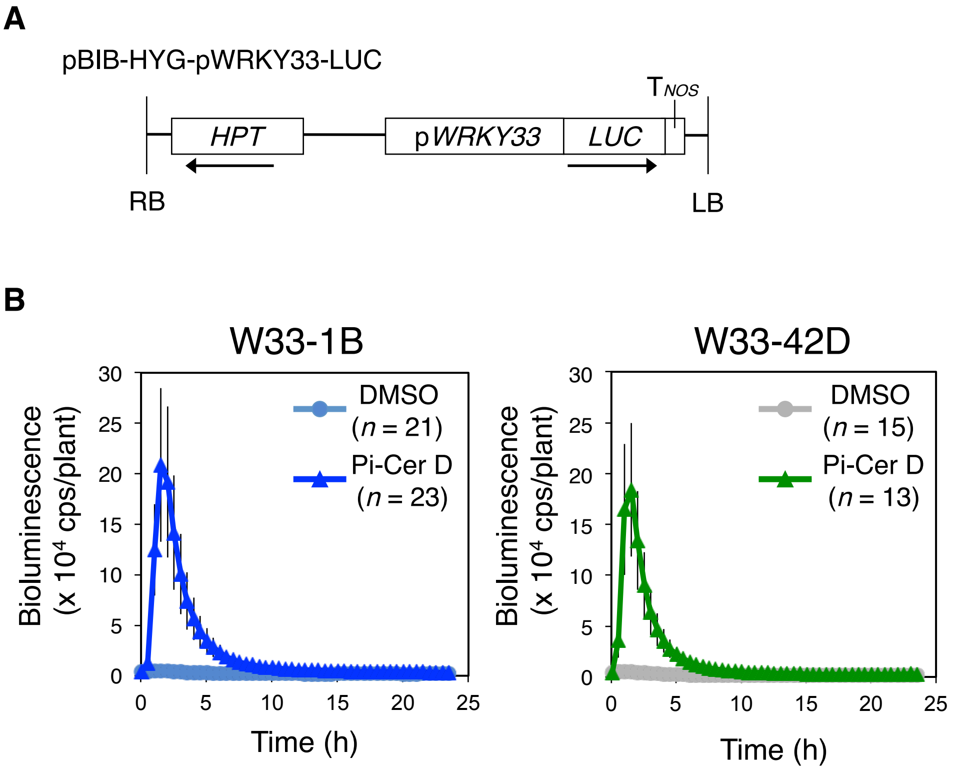


**Fig. S2. p*WRKY33-LUC* reporter line.** (A) Structure of the p*WRKY33-LUC* reporter construct: *HPT*, a hygromycin resistance gene cassette in pBIB-HYG; p*WRKY33*, the promoter of *WRKY33* (at positions -2,000 to -1); *LUC*, the coding region of a modified *luciferase* gene derived from *Photinus pyralis*; $T_{NOS}$, transcriptional terminator of *NOS* (*nopaline synthase*) from *Agrobacterium*; LB, left-border sequence of T-DNA; RB, right-border sequence of T-DNA. The direction of transcription of the gene cassettes is shown by arrows. (B) Luciferase-mediated bioluminescence patterns of p*WRKY33-LUC* reporter lines. Eight-day-old seedlings of W33-1B and W33-42D were treated with water or 0.17 μM Pi-Cer D. Bioluminescence from each seedling was monitored with a real-time bioluminescence monitoring system at the indicated time points. Data are means ± SD.

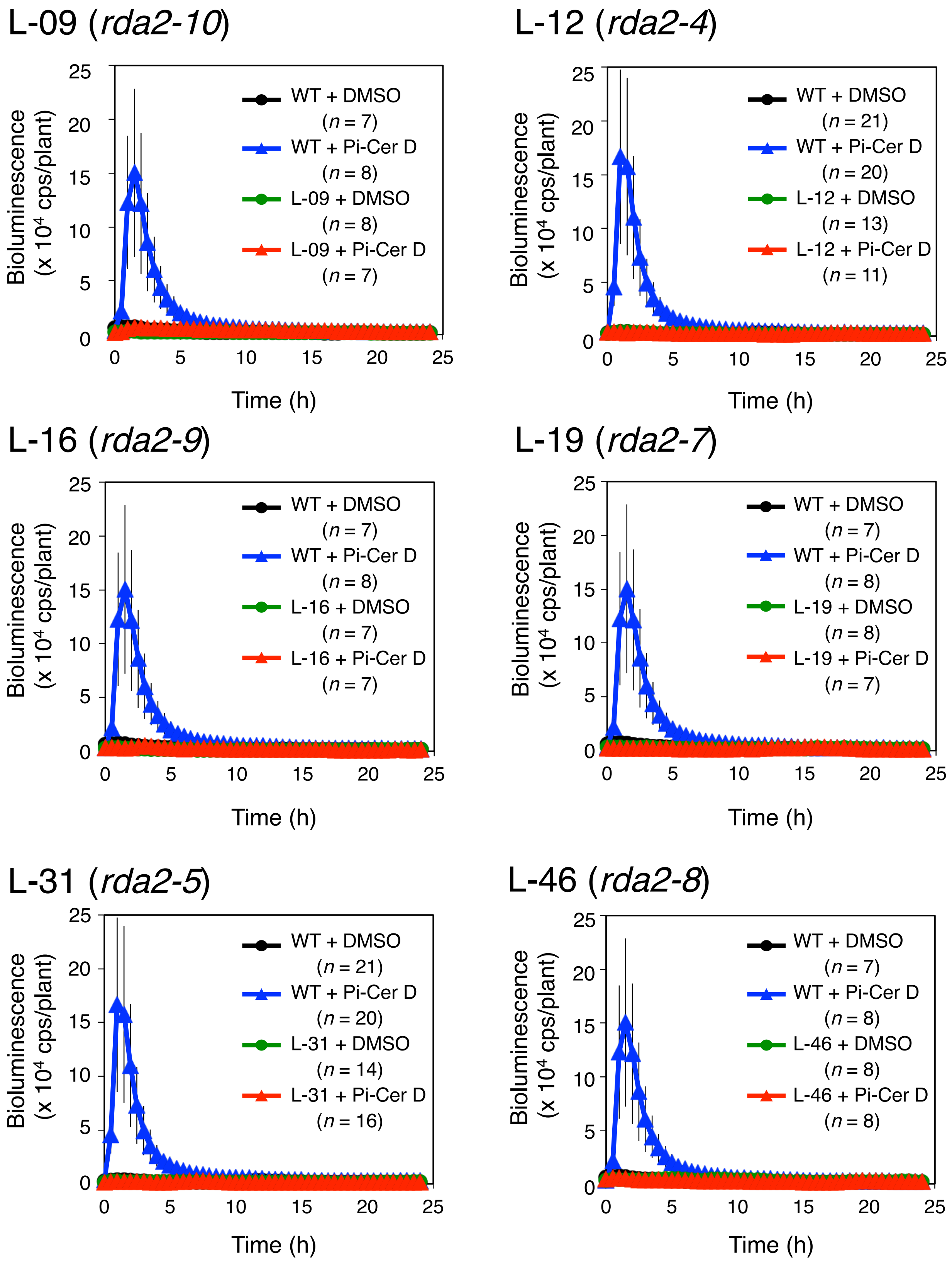
L-09 (rda2-10)
WT + DMSO (n = 7)
WT + Pi-Cer D (n = 8)
L-09 + DMSO (n = 8)
L-09 + Pi-Cer D (n = 7)
L-12 (rda2-4)
WT + DMSO (n = 21)
WT + Pi-Cer D (n = 20)
L-12 + DMSO (n = 13)
L-12 + Pi-Cer D (n = 11)
L-16 (rda2-9)
WT + DMSO (n = 7)
WT + Pi-Cer D (n = 8)
L-16 + DMSO (n = 7)
L-16 + Pi-Cer D (n = 7)
L-19 (rda2-7)
WT + DMSO (n = 7)
WT + Pi-Cer D (n = 8)
L-19 + DMSO (n = 8)
L-19 + Pi-Cer D (n = 7)
L-31 (rda2-5)
WT + DMSO (n = 21)
WT + Pi-Cer D (n = 20)
L-31 + DMSO (n = 14)
L-31 + Pi-Cer D (n = 16)
L-46 (rda2-8)
WT + DMSO (n = 7)
WT + Pi-Cer D (n = 8)
L-46 + DMSO (n = 8)
L-46 + Pi-Cer D (n = 8)
Bioluminescence (x 10⁴ cps/plant)
Time (h)
0 5 10 15 20 25

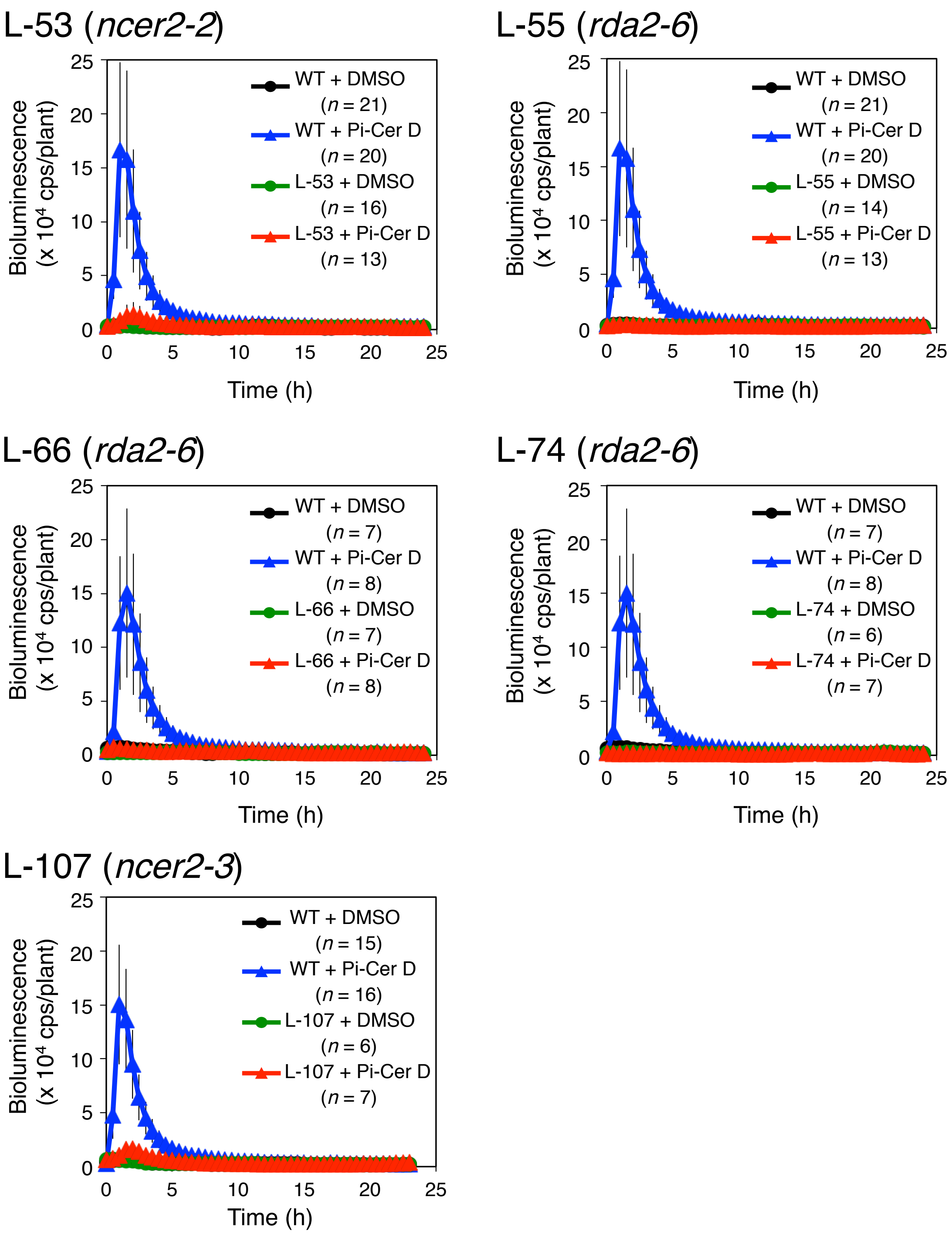


**Fig. S3. Bioluminescence patterns of mutants with altered responses to Pi-Cer D treatment.** Eight-day-old seedlings of p*WRKY33-LUC* reporter (WT) and mutant lines treated with DMSO (0.005%) or 0.17 μM Pi-Cer D. Bioluminescence from each seedling was monitored with a real-time bioluminescence monitoring system at the indicated time points. Data are means ± SD. Experiments were conducted three times with similar results.

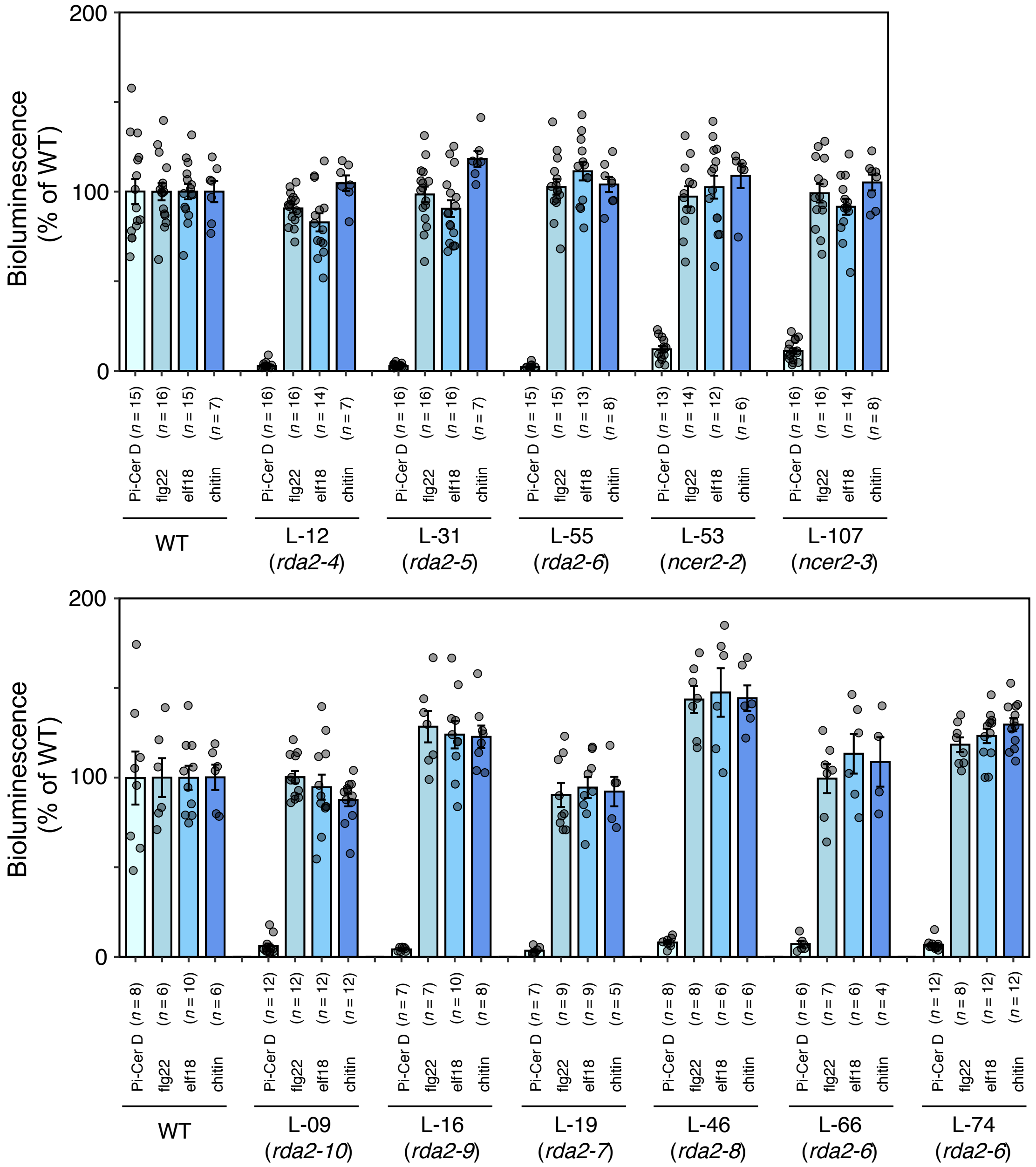


**Fig. S4. Bioluminescence responses to different PAMPs of 11 mutants with altered responses to Pi-Cer D treatment.** Eight-day-old p*WRKY33-LUC* reporter (WT) and mutant seedlings were treated with 0.17 µM Pi-Cer D, 0.5 µM flg22, 0.5 µM elf18 or 20 µg mL$^{-1}$ chitin. Response to the four elicitors was monitored with a real-time bioluminescence monitoring system. Bioluminescence is shown as % of WT. Data are peak means ± SE.

### L-09 (*rda2-10*)

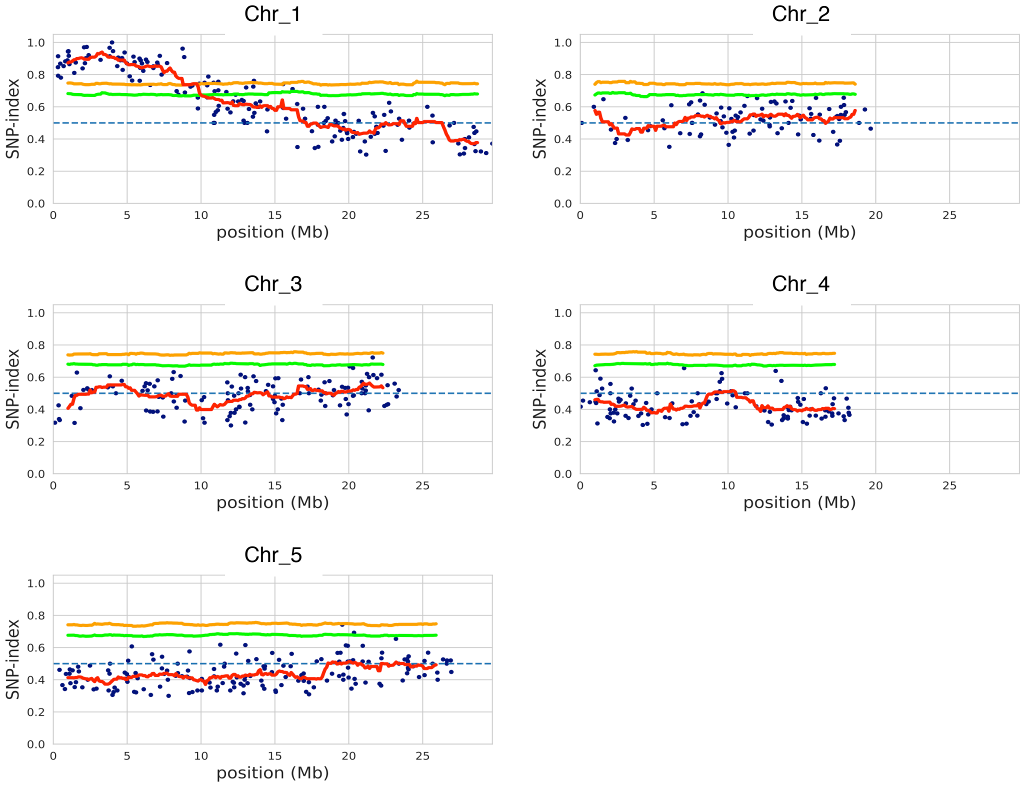


### L-12 (*rda2-4*)

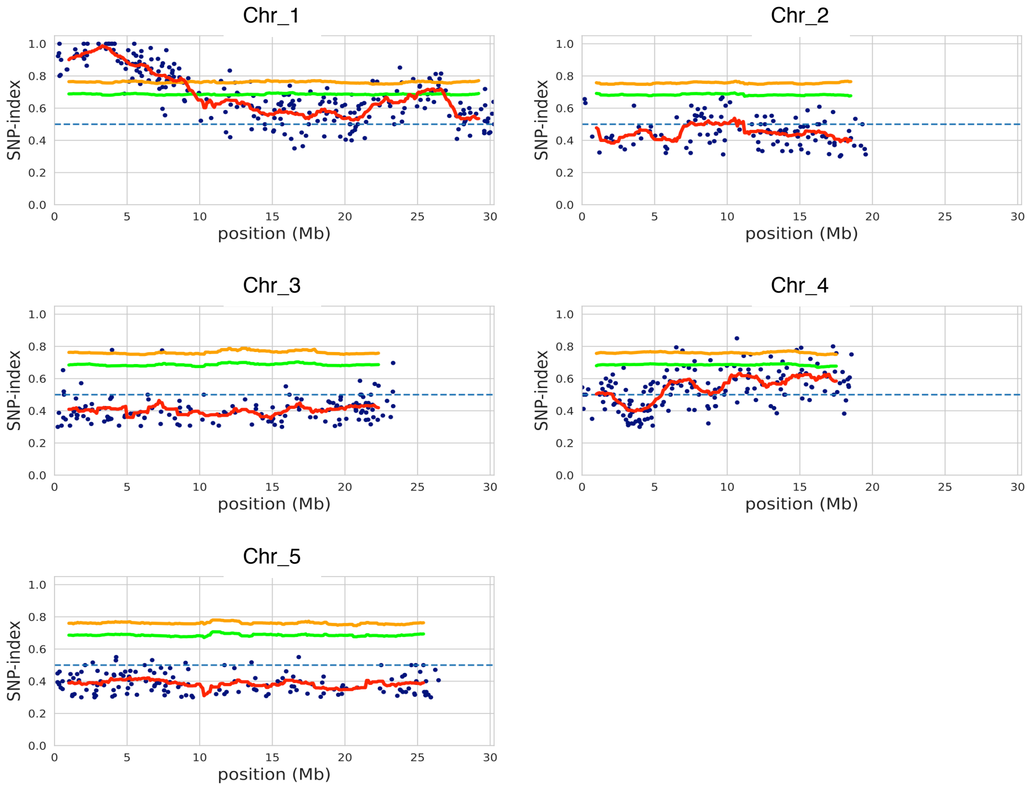

### L-16 (*rda2-9*)

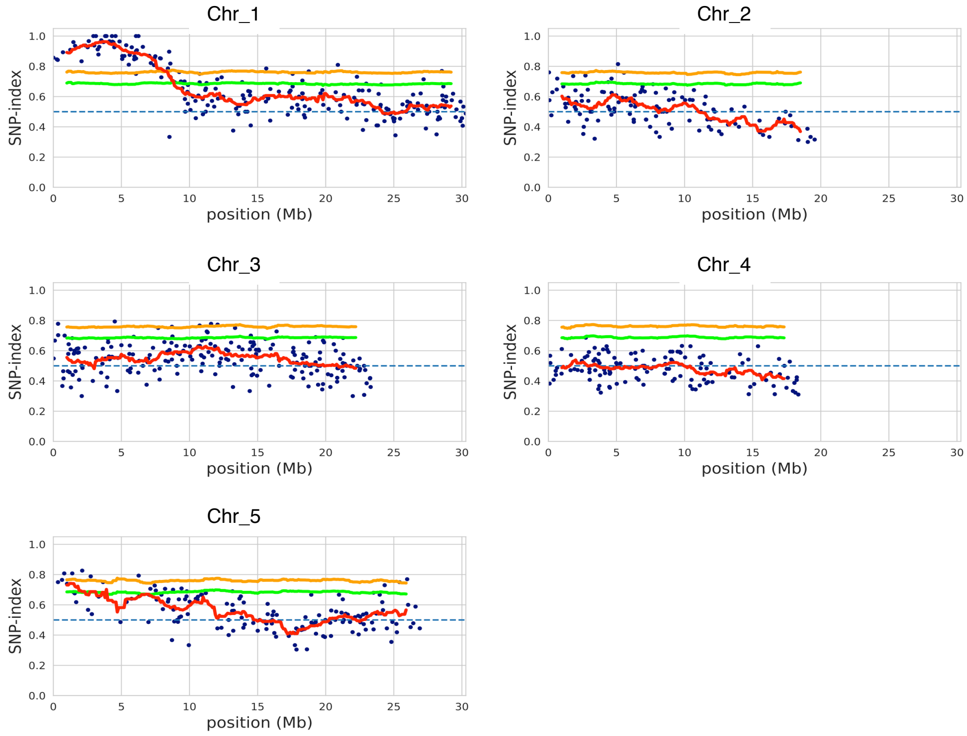


### L-19 (*rda2-7*)

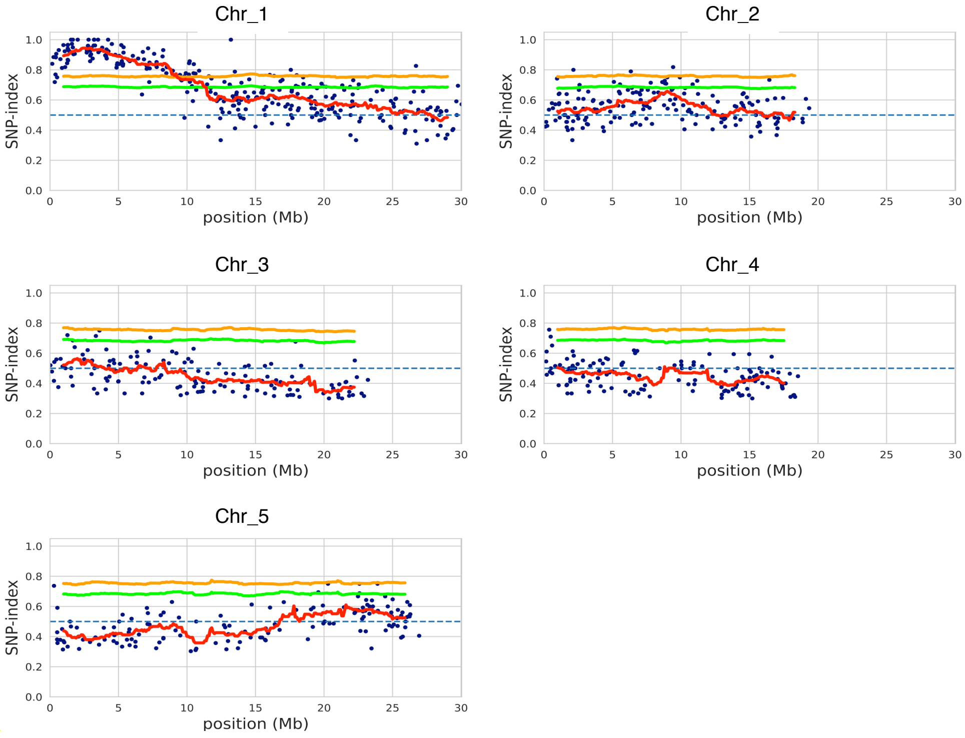

### L-31 (*rda2-5*)

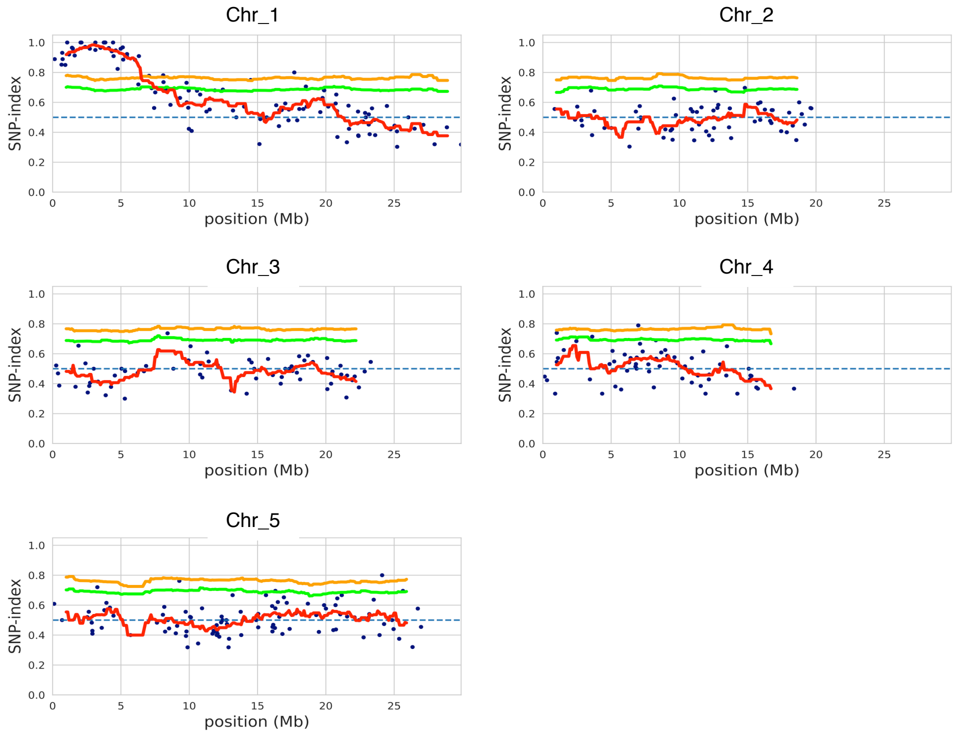


### L-46 (*rda2-8*)

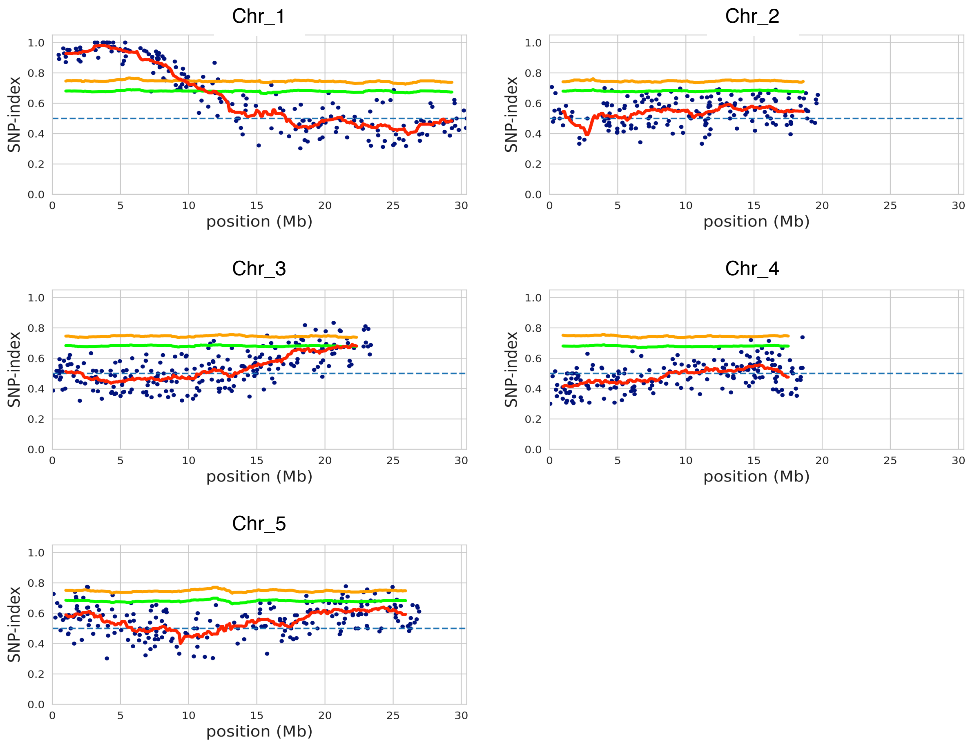

### L-55 (*rda2-6*)

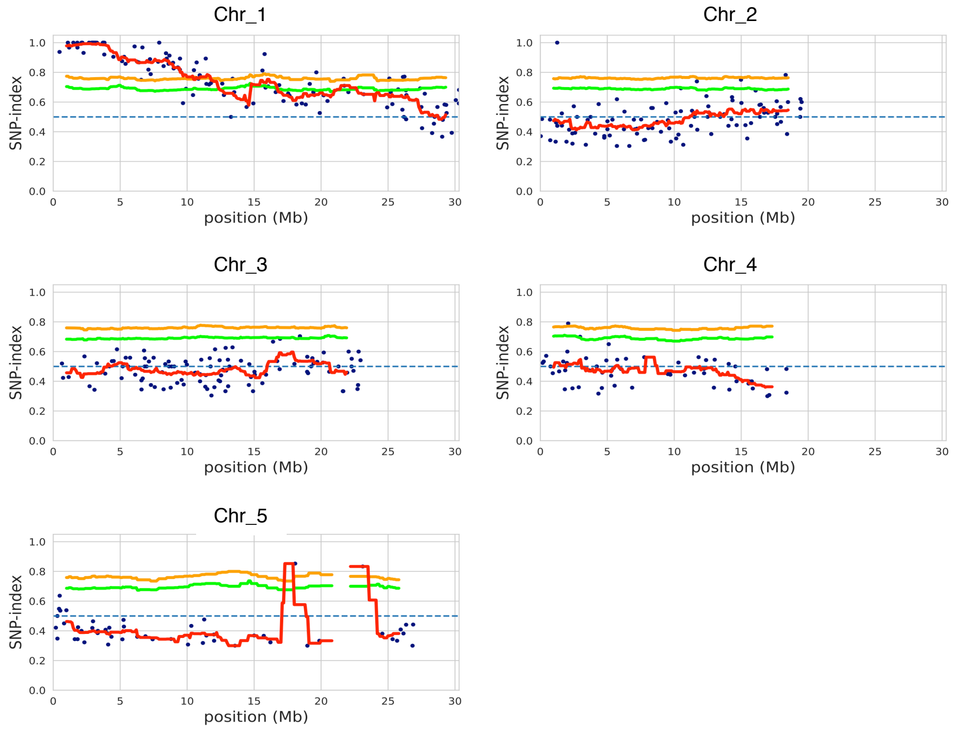


### L-66 (*rda2-6*)

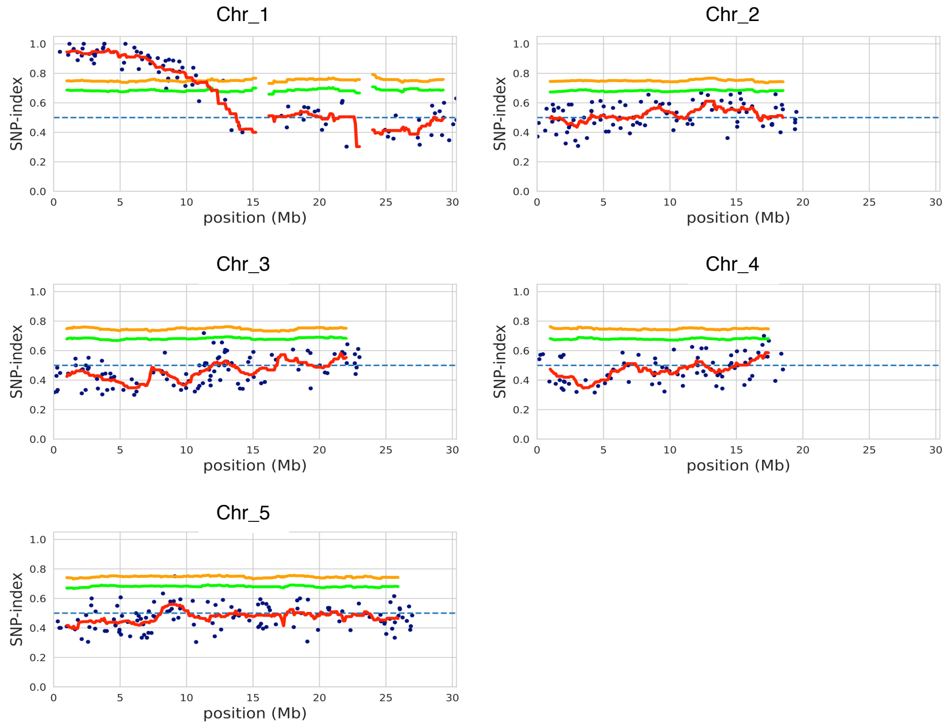

#### L-74 (*rda2-6*)

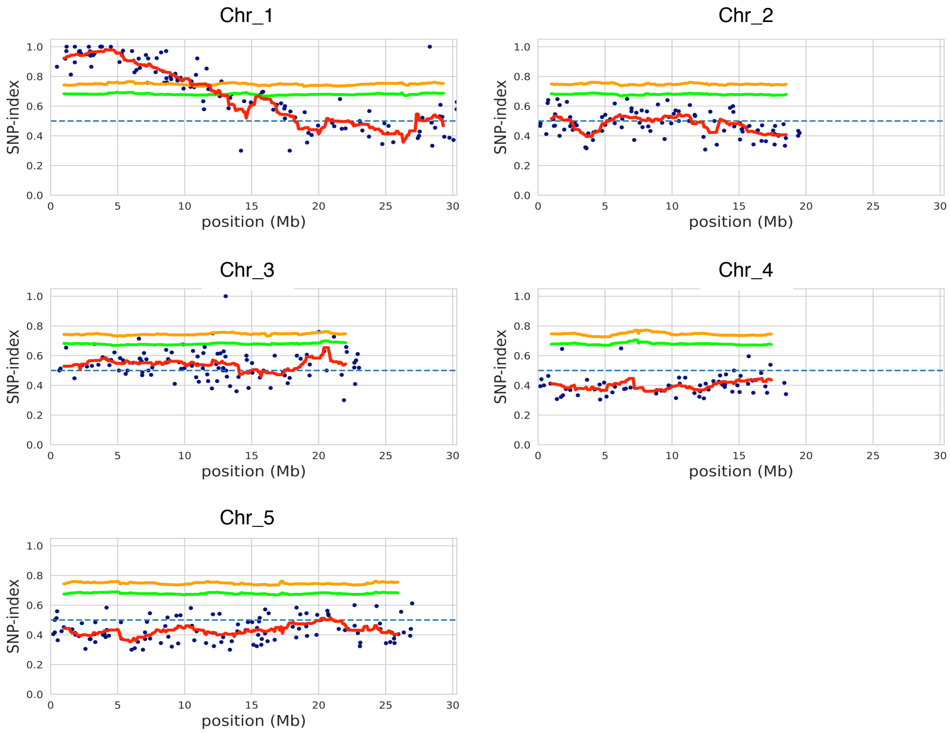


#### L-53 (*ncer2-2*)

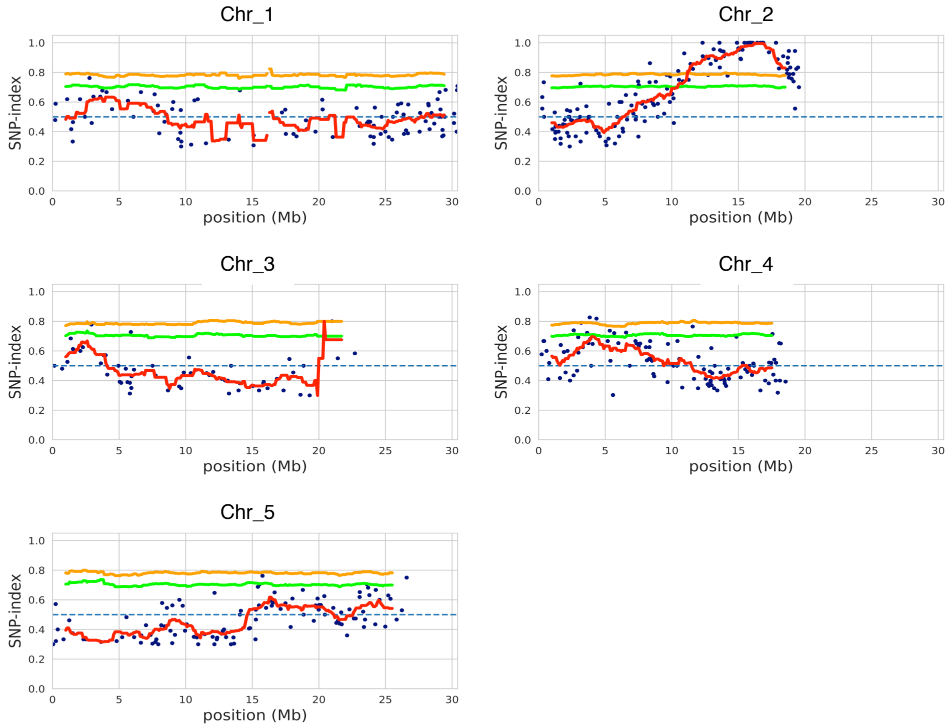

#### L-107 (*ncer2-3*)

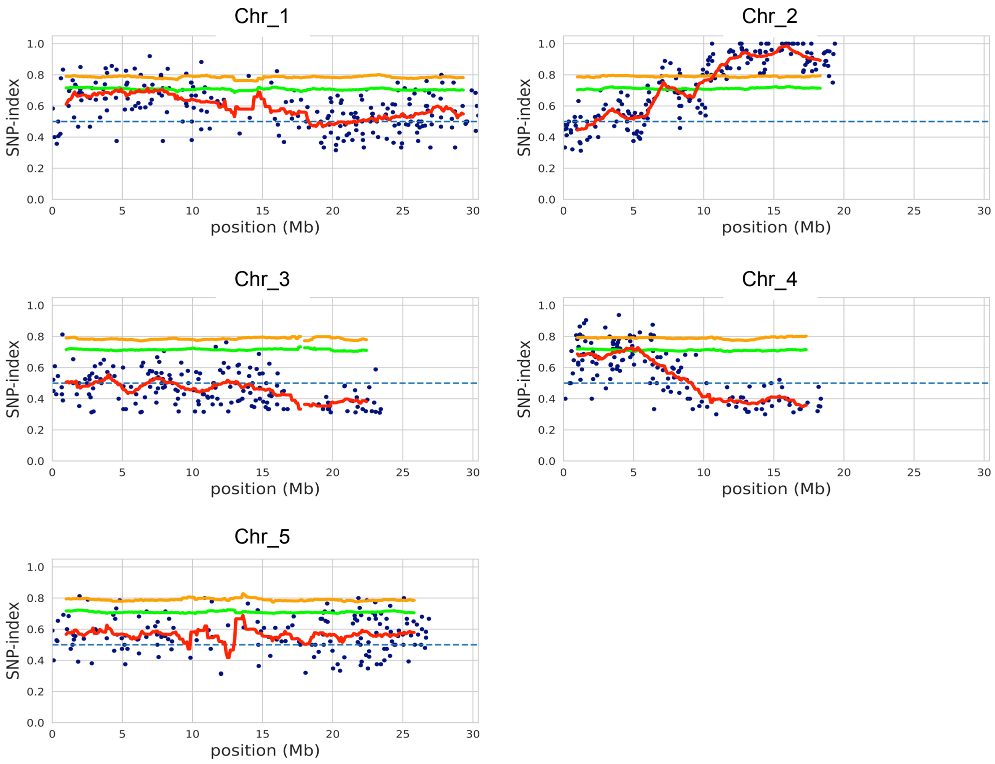


**Fig. S5. MutMap SNP-index plots for 11 Arabidopsis mutants with altered responses to Pi-Cer D treatment.** MutMap SNP-index plots of the five Arabidopsis chromosomes. The genomic region with the highest SNP-index peak indicates the location of the causal mutation. Each mutant was crossed with the W33-1B line and the resulting $F_2$ progeny were tested for bioluminescence. The DNA from 30 $F_2$ progeny with mutant phenotypes was bulked and subjected to whole-genome sequencing and MutMap analysis. Blue dots represent SNPs in the mutant. The red line represents mean SNP-index values across a 2-Mb sliding window with 10-kb increments. The green and yellow lines show the 95% or 99% confidence limit, respectively, of SNP-index values under the null hypothesis of SNP-index = 0.5.

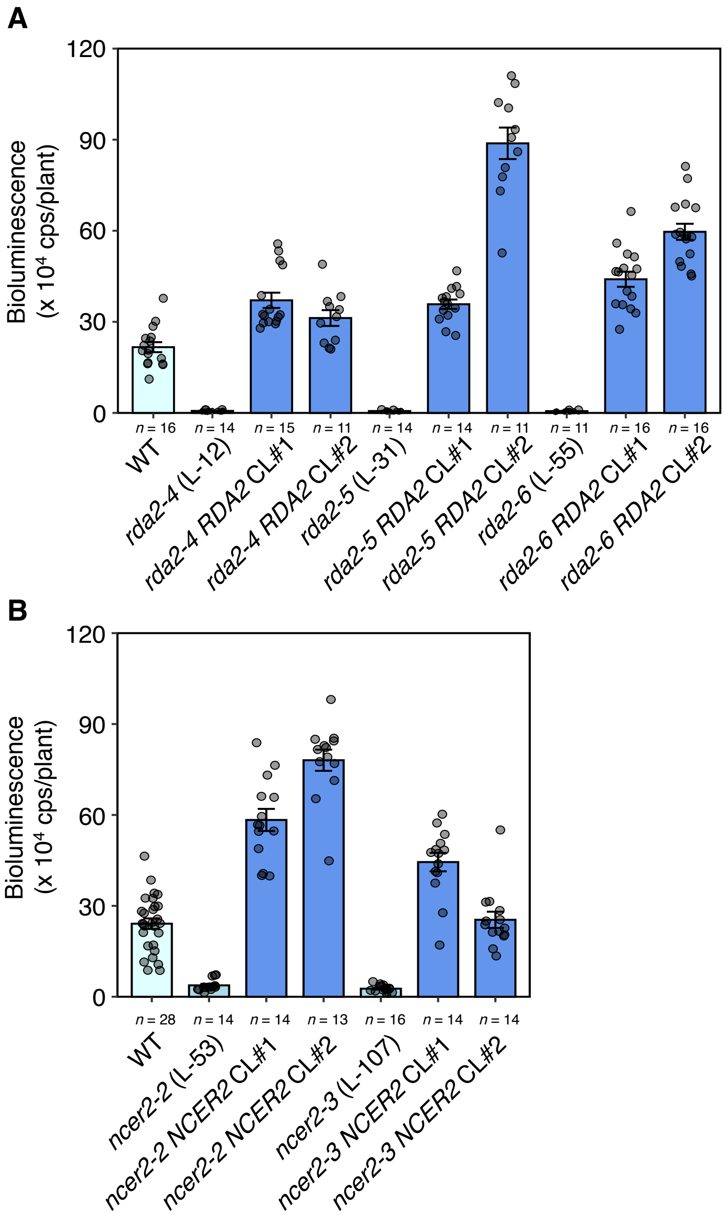


**Fig. S6. Genetic complementation of *rda2* and *ncer2* mutants.** (A) Bioluminescence analysis of the *rda2 RDA2* complementation lines (CL). (B) Bioluminescence analysis of the *ncer2 NCER2* complementation lines. Eight-day-old seedlings of p*WRKY33-LUC* reporter (WT), *rda2*, *ncer2*, and each complementation line were treated with 0.17 μM Pi-Cer D. Peak bioluminescence values are shown (means ± SE).

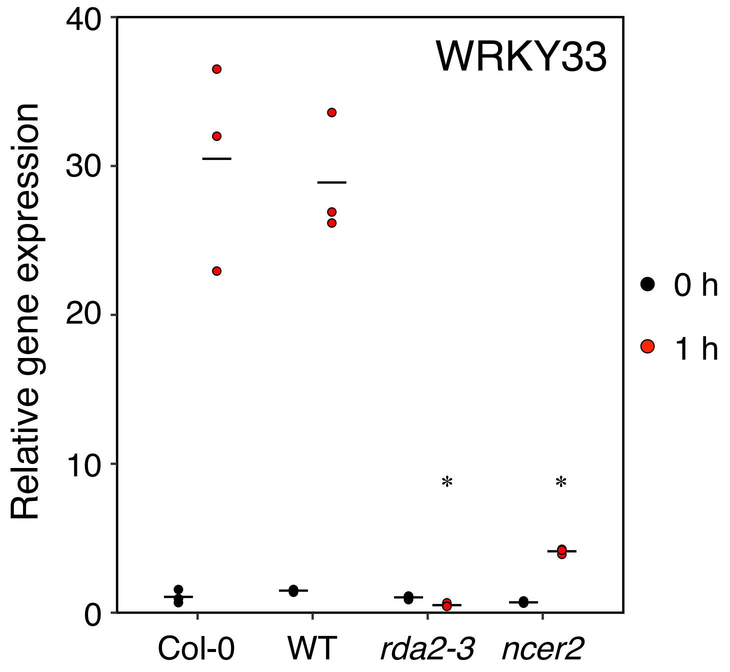


**Fig. S7. *WRKY33* gene expression in *rda2* and *ncer2* T-DNA insertion mutant lines.** Eight-day-old seedlings of Col-0, p*WRKY33-LUC* reporter (WT), *rda2-3*, and *ncer2* were treated with 0.17 μM Pi-Cer D. The expression of *WRKY33* is relative to that of the *UBC* housekeeping gene and was normalized to values for Col-0 seedlings (0 h). Individual data (symbols) and means (bars) are shown ($n$ = 3). *, $p < 0.05$ in two-tailed $t$-tests compared with the corresponding values for Col-0 at each time point.

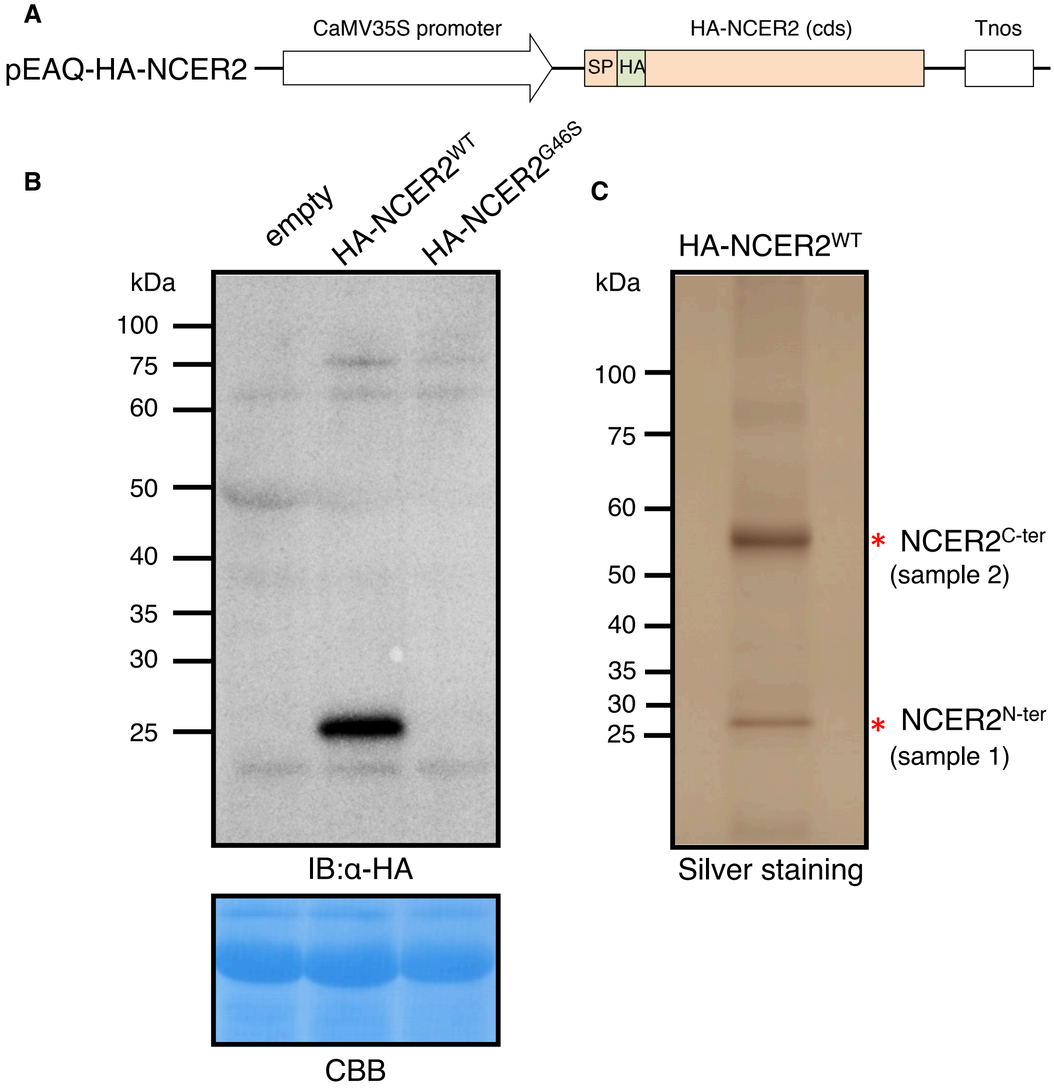


**Fig. S8. NCER2 is cleaved into N- and C-terminal fragments that form a complex.** (A) Schematic representation of the HA-NCER2 construct expressed in *Nicotiana benthamiana*. (B) Anti-HA immunoblot analysis of HA-NCER2 expressed in *N. benthamiana*. A CBB-stained loading control is shown (bottom). (C) Silver-stained image of the HA-purified fraction from *N. benthamiana* expressing HA-NCER2. The smaller (sample 1) and larger (sample 2) bands were identified by MS analysis as the C-and N-terminal regions of NCER2, respectively.

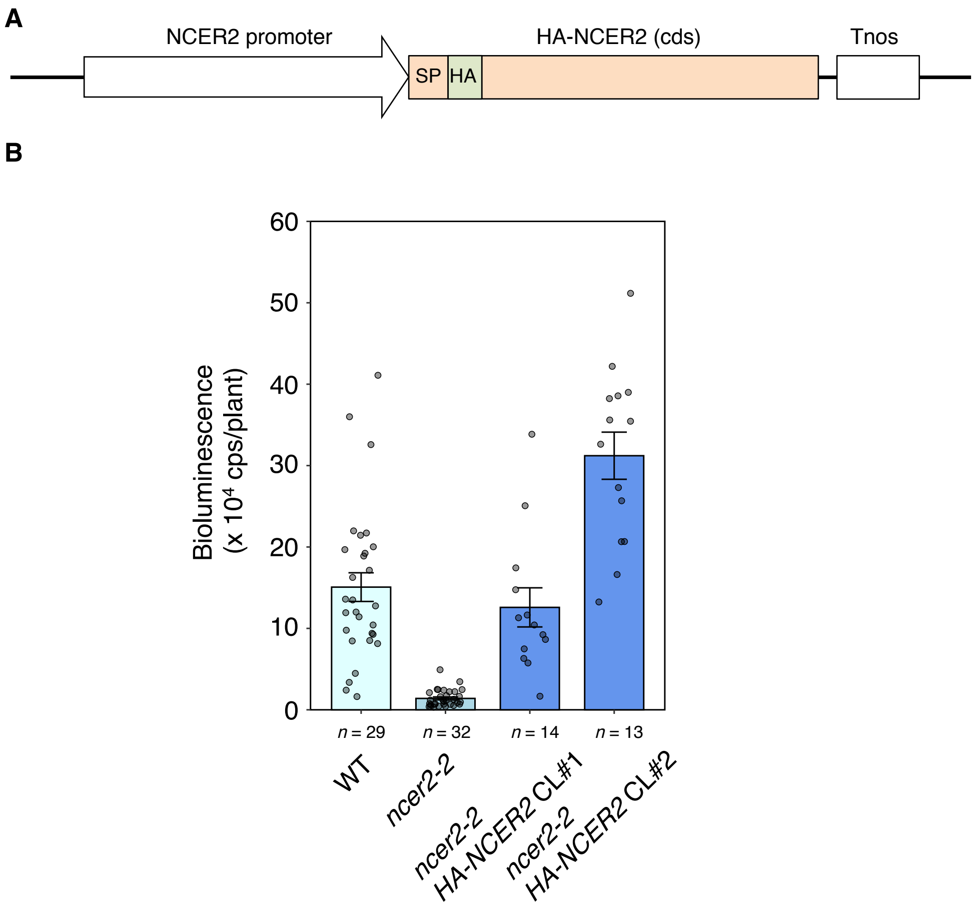


**Fig. S9. Genetic complementation of the *ncer2* mutant with HA-NCER2.** (A) Schematic representation of the HA-NCER2 construct expressed in the Arabidopsis *ncer2-2* mutant. (B) Bioluminescence analysis of the *ncer2-2 HA-NCER2* complementation lines (CL). Eight-day-old seedlings of p*WRKY33-LUC* reporter (WT), *ncer2-2* and *ncer2-2 HA-NCER2* complementation lines were treated with 0.17 μM Pi-Cer D. Peak bioluminescence values are shown (means ± SE).

**A**

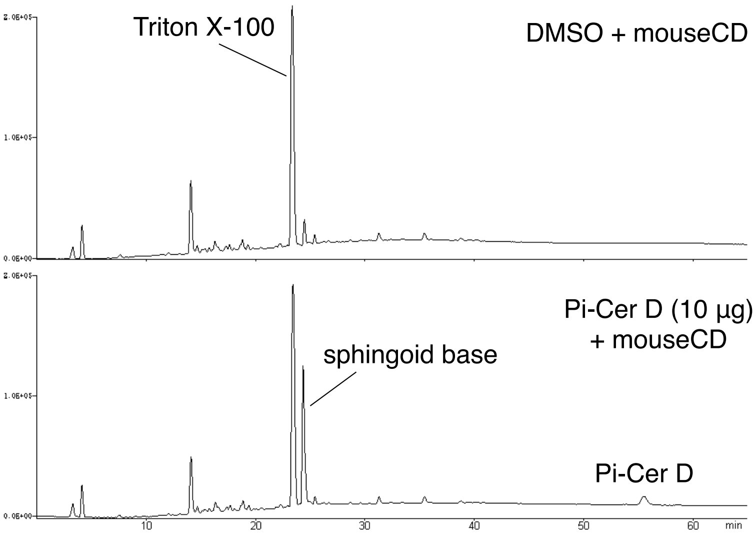


**B**

Reaction mixture from Pi-Cer D (10 µg) + mouseCD

dissolved in MeOH (100 µL)

10 mL (corresponding to 1 µg of P-iCer D)

HPLC: Develosil ODS-UG-5 (4.6 x 250 mm),
90-100% A in 20 mM $NH_4OAc$ (10 min),
A = MeOH-2 M $NH_4OAc$ (99:1), 1 mL/min, 245 nm

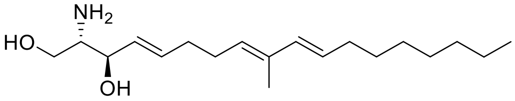


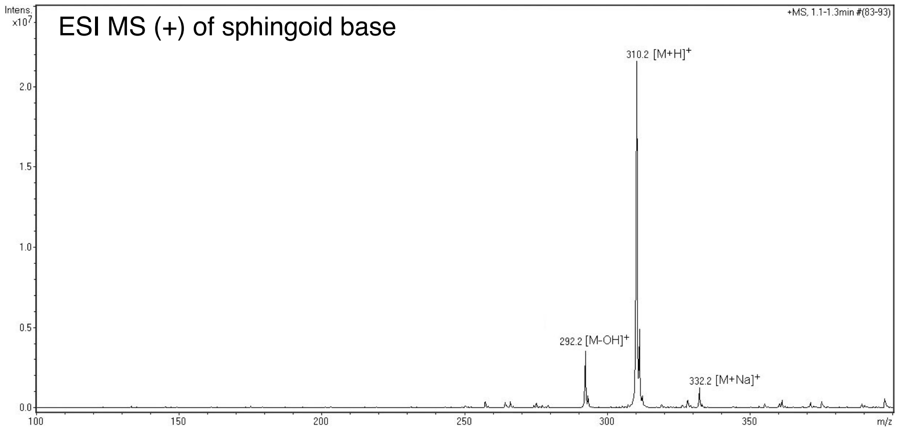


**Fig. S10. Detection and purification of sphingoid base cleaved from Pi-Cer D.** (A) HPLC analysis of the enzymatic reaction mixture. (B) Purification protocol and mass spectrum (ESI MS, positive ion) of sphingoid base from the reaction mixture of Pi-Cer D with mouse CD.

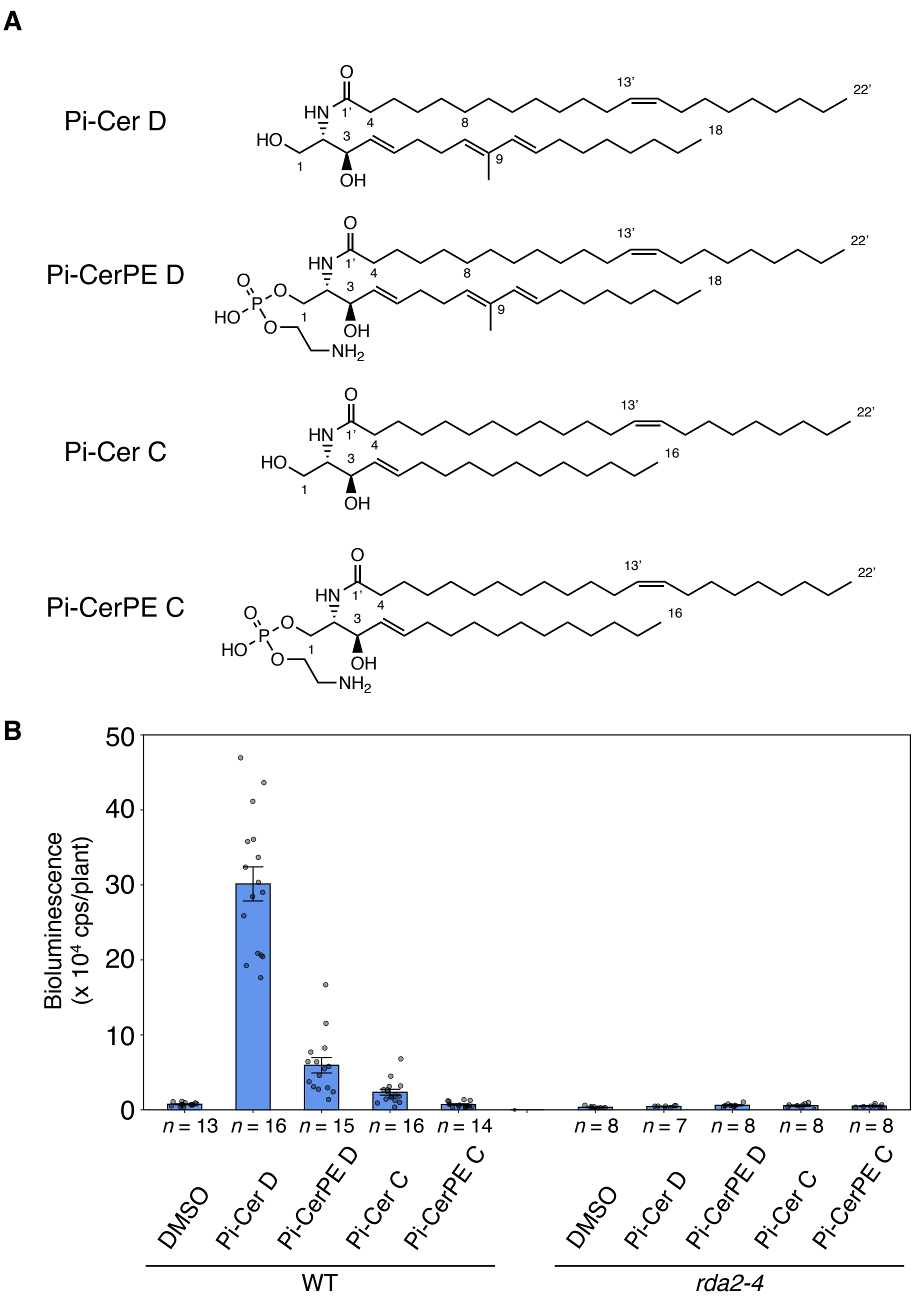


**Fig. S11. Differences in sphingoid base structures of Pi-Cer D-derivatives affect the induction of defense response.** (A) Structures of Pi-Cer D and its derivatives. (B) Bioluminescence analysis for the p*WRKY33-LUC* reporter (WT) and *rda2-4* mutant treated with Pi-Cer D and its derivatives. Eight-day-old seedlings were treated with Pi-Cer D, Pi-CerPE D, Pi-Cer C, or Pi-CerPE C (0.17 µM). Peak bioluminescence values are shown (means ± SE).

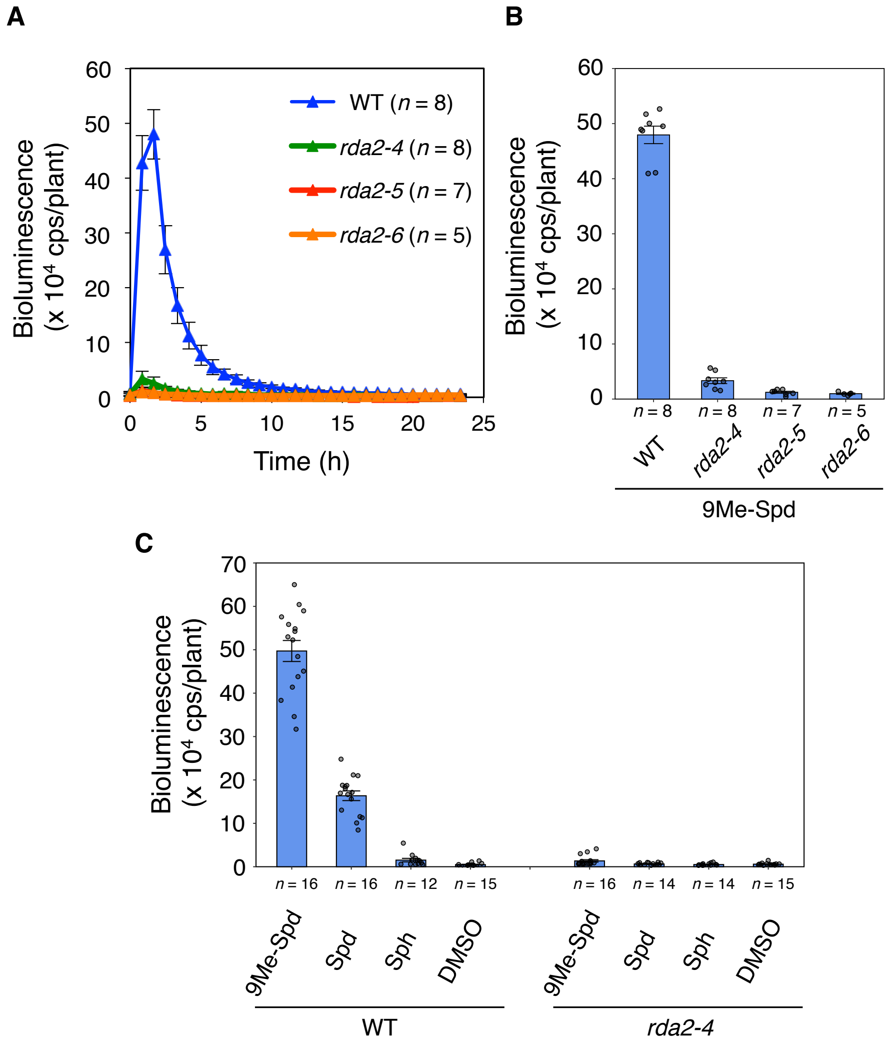


**Fig. S12. RDA2-dependent bioluminescence induction by treatment with sphingoid bases.** (A and B) Luciferase-mediated bioluminescence patterns (A) and peak bioluminescence (B) for the p*WRKY33-LUC* reporter (WT) and *rda2* mutants treated with (4*E*,8*E*)-9-methyl-4,8-sphingadienine (9Me-Spd) (0.5 µM). (C) Bioluminescence analysis for WT and *rda2-4* mutant treated with sphingoid bases. Eight-day-old seedlings were treated with 9Me-Spd, (4*E*,8*E*)-4,8-sphingadienine (Spd), or Sphingosine (Sph) (0.5 µM). (B and C) Peak bioluminescence values are shown (means ± SE).

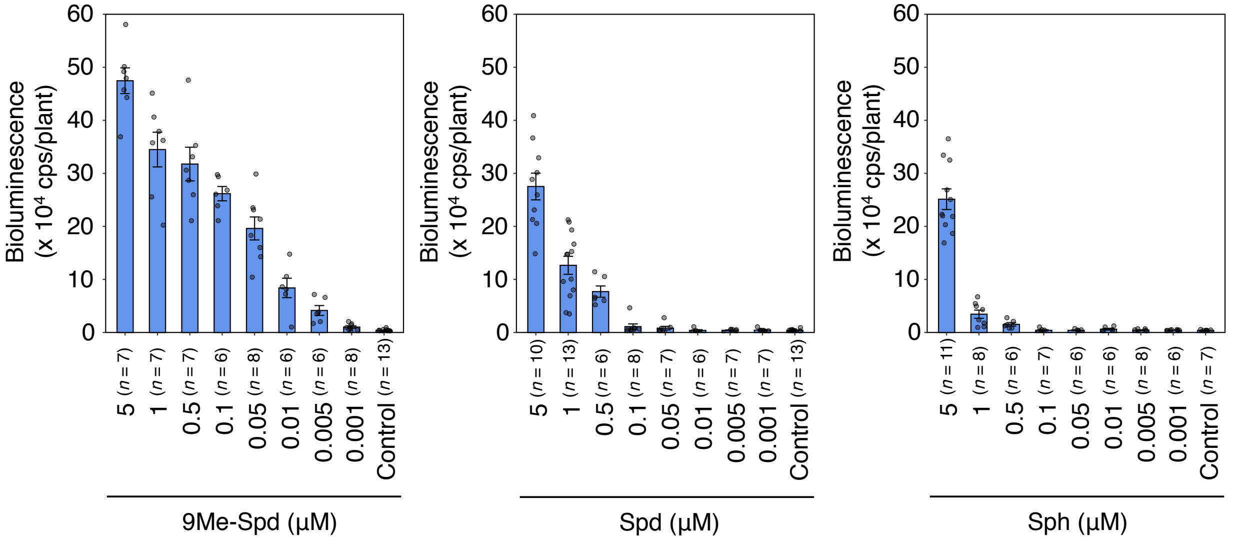


**Fig. S13. Concentration-dependent bioluminescence response of Arabidopsis seedlings to sphingoid bases.** Eight-day-old p*WRKY33-LUC* reporter seedlings were treated with (4*E*,8*E*)-9-methyl-4,8-sphingadienine (9Me-Spd), (4*E*,8*E*)-4,8-sphingadienine (Spd), or sphingosine (Sph) in different concentrations. Peak bioluminescence values are shown (means ± SE).

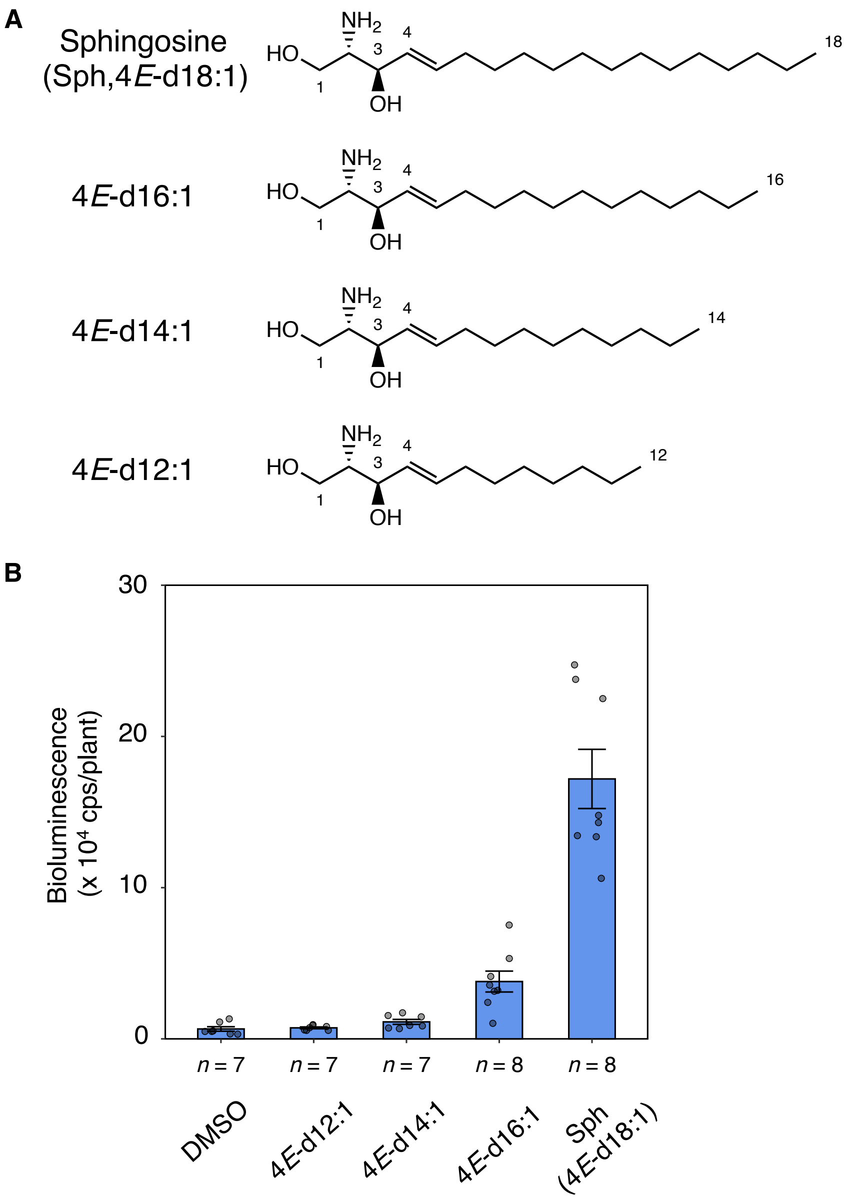


**Fig. S14. Carbon chain length in sphingosine affects defense induction activity.** (A) Structures of sphingosine (4*E*-d18:1, Sph) and its derivatives. (B) Bioluminescence analysis for the p*WRKY33-LUC* reporter treated with sphingoid bases. Eight-day-old seedlings were treated with 4*E*-d12:1, 4*E*-d14:1, 4*E*-d16:1, or Sph (5 μM). Peak bioluminescence values are shown (means ± SE).

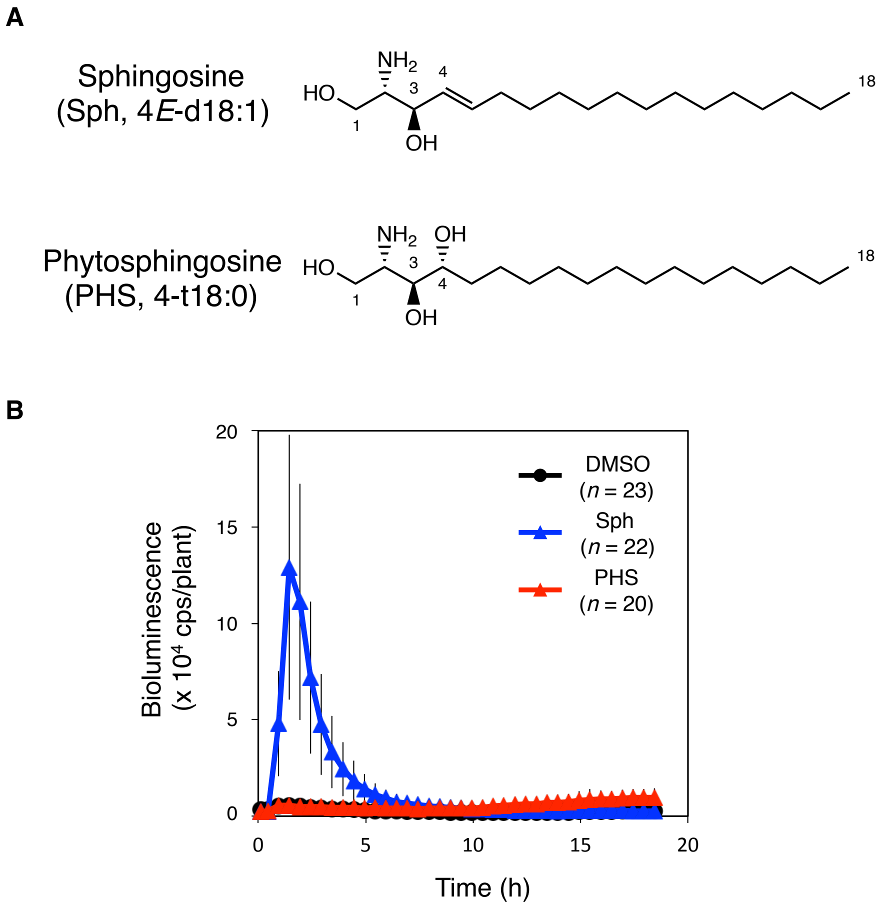


**Fig. S15. The 4-double bond structure in sphingoid base affects defense induction.** (A) Structures of sphingosine and its derivatives. (B) Bioluminescence analysis for the p*WRKY33-LUC* reporter (WT) treated with sphingoid bases. Eight-day-old seedlings were treated with sphingosine (Sph, 5 μM) or phytosphingosine (PHS, 5 μM). Data are means ± SD.

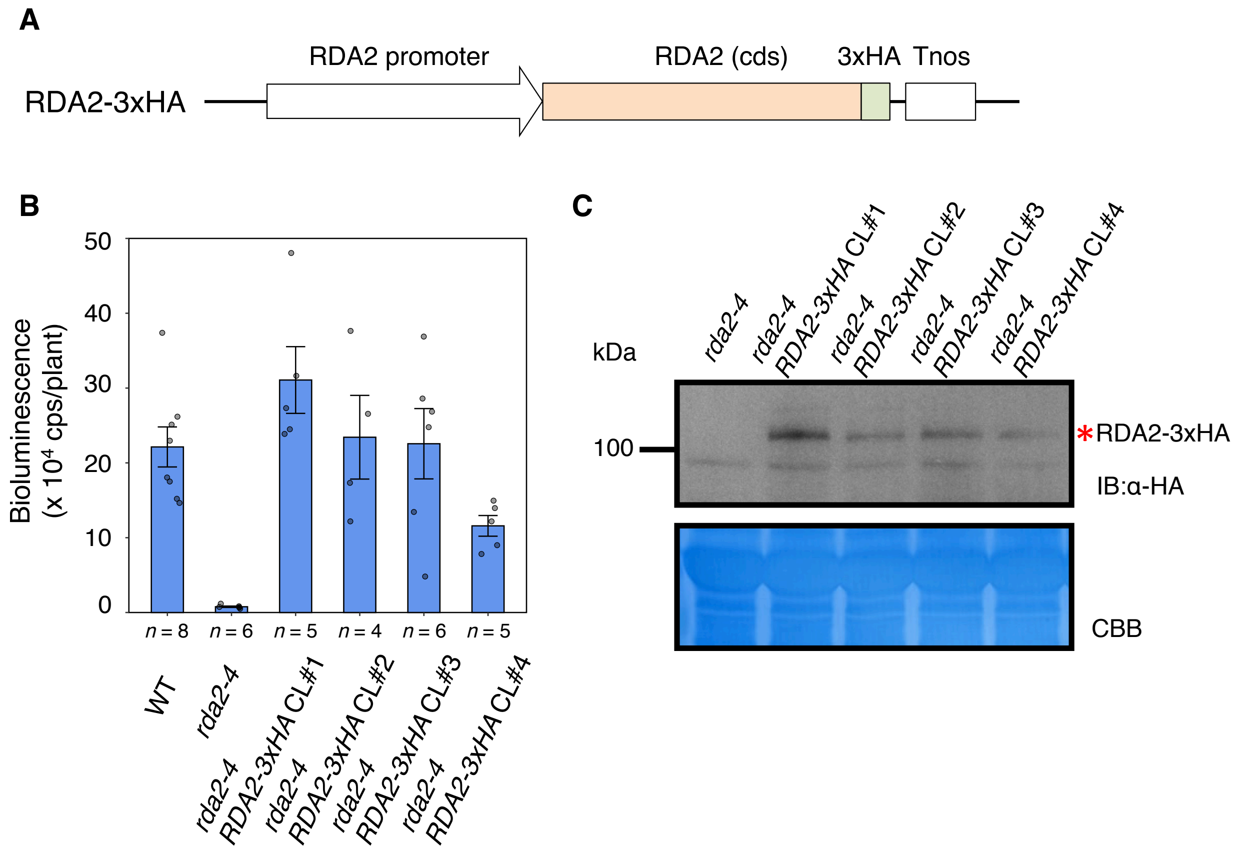


**Fig. S16. RDA2-3×HA is a functional protein.** (A) Schematic representation of the RDA2-3×HA construct expressed in Arabidopsis *rda2-4*. (B) Bioluminescence analysis of *rda2-4 RDA2-3×HA* complementation lines. Eight-day-old seedlings of the p*WRKY33-LUC* reporter (WT), *rda2-4*, and *rda2-4 RDA2-3×HA* complementation lines (CL) were treated with 0.17 μM Pi-Cer D. Peak bioluminescence values are shown (means ± SE). (C) Anti-HA immunoblot analysis for RDA2-3×HA expressed in the Arabidopsis *rda2-4* mutant. A red asterisk indicate RDA2-3×HA. A CBB-stained loading control is shown below.

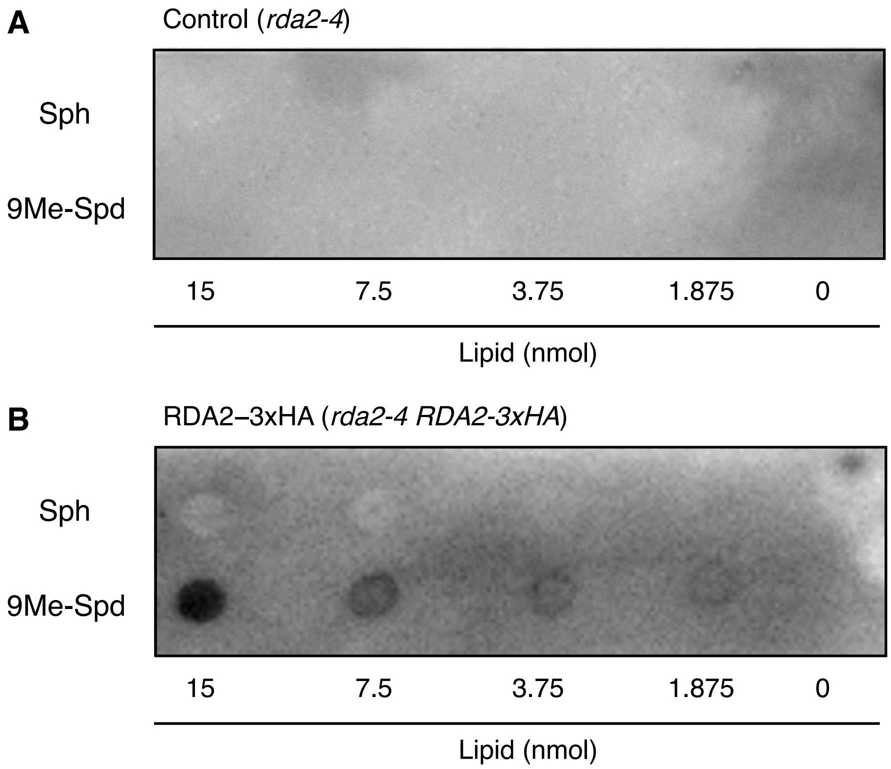


**Fig. S17. Interaction between sphingoid bases and RDA2 demonstrated by protein-lipid overlay assay.** Sphingosine (Sph) and (4*E*,8*E*)-9-methyl-4,8-sphingadienine (9Me-Spd) were spotted onto a PVDF membrane in the indicated amounts. Membrane protein fractions were prepared from the Arabidopsis *rda2-4* mutant (A) or *rda2-4 RDA2-3×HA* plants (B). The assays were performed simultaneously using the same ECL reagent and settings.

**Table S1. List of mutants used in this study.**

| Allele name | Mutant no. | Accession no. | Mutation | Background |
|---|---|---|---|---|
| *rda2-10* | L-09 | At1g11330 | G841R | p*WRKY33-LUC* reporter (Col-0) |
| *rda2-4* | L-12 | At1g11330 | R220H | p*WRKY33-LUC* reporter (Col-0) |
| *rda2-9* | L-16 | At1g11330 | G533S | p*WRKY33-LUC* reporter (Col-0) |
| *rda2-7* | L-19 | At1g11330 | A109T | p*WRKY33-LUC* reporter (Col-0) |
| *rda2-5* | L-31 | At1g11330 | E700K | p*WRKY33-LUC* reporter (Col-0) |
| *rda2-8* | L-46 | At1g11330 | G222R | p*WRKY33-LUC* reporter (Col-0) |
| *rda2-6* | L-55 | At1g11330 | G585R | p*WRKY33-LUC* reporter (Col-0) |
| *rda2-6* | L-66 | At1g11330 | G585R | p*WRKY33-LUC* reporter (Col-0) |
| *rda2-6* | L-74 | At1g11330 | G585R | p*WRKY33-LUC* reporter (Col-0) |
| *rda2-3* | SALK_143489C | At1g11330 | T-DNA insertion (chr1_3812038-3812103) | Col-0 |
| *ncer2-2* | L-53 | At2g38010 | Q88* | p*WRKY33-LUC* reporter (Col-0) |
| *ncer2-3* | L-107 | At2g38010 | G46S | p*WRKY33-LUC* reporter (Col-0) |
| *ncer2* | SALK_050305 | At2g38010 | T-DNA insertion (chr2_15907314-15907352) | Col-0 |

#### Table S2. Candidate genes for causal mutations identified by MutMap.

| Mutant lines | Chromosome coordinate | Reference base | Altered base | Depth | SNP-index | Accession | Note | Type of mutation |
|---|---|---|---|---|---|---|---|---|
| L-09 *(rda2-10)* | 3813408 | G | A | 28 | 0.96 | AT1G11330 | S-locus lectin protein kinase family protein | G841R |
| | 4045211 | G | A | 33 | 0.93 | AT1G11980 | ubiquitin-related protein 3 | -409 |
| | 4736600 | G | A | 43 | 0.9 | AT1G13820 | alpha/beta-Hydrolases superfamily protein | A162T |
| | 4827272 | G | A | 31 | 0.96 | AT1G14100 | fucosyltransferase 8 | -690 |
| L-12 *(rda2-4)* | 3033029 | G | A | 23 | 1 | AT1G09390 | GDSL-like Lipase/Acylhydrolase superfamily protein | intron |
| | 3136805 | G | A | 33 | 0.96 | AT1G09690 | Translation protein SH3-like family protein | intron |
| | 3554807 | G | A | 31 | 0.96 | AT1G10700 | phosphoribosyl pyrophosphate synthase 3 | intron |
| | 3555640 | G | A | 27 | 1 | AT1G10700 | phosphoribosyl pyrophosphate synthase 3 | intron |
| | 3705317 | G | A | 22 | 1 | AT1G11100 | SNF2-RING-HELICASE?LIKE 5 | A878V |
| | 3710954 | G | A | 26 | 1 | AT1G11110 | LisH and RanBPM domains containing protein | V104I |
| | 3811030 | G | A | 18 | 1 | AT1G11330 | S-locus lectin protein kinase family protein | R220H |
| | 3950098 | G | A | 28 | 1 | AT1G11710 | putative salt-inducible protein | G405S |
| | 3961816 | G | A | 26 | 0.96 | AT1G11735 | MIR171b | -428 |
| | 4127150 | G | A | 27 | 1 | AT1G12160 | Flavin-binding monooxygenase family protein | A273T |
| | 4134396 | G | A | 29 | 0.96 | AT1G12190 | F-box and associated interaction domains-containing protein | -301 |
| | 4360527 | G | A | 27 | 0.92 | AT1G12790 | DNA ligase-like protein | intron |
| L-16 (*rda2-9*) | 3004160 | G | A | 30 | 0.93 | AT1G09300 | Metallopeptidase M24 family protein | intron |
| | 3126271 | G | A | 18 | 1 | AT1G09650 | F-box and associated interaction domains-containing protein | W98* |
| | 3261888 | G | A | 31 | 0.93 | AT1G09980 | Putative serine esterase family protein | -1021 |
| | 3372864 | G | A | 27 | 0.92 | AT1G10290 | dynamin-like protein 6 | G308E |
| | 3686297 | G | A | 20 | 0.95 | AT1G11060 | AtWAPL1 | intron |
| | 3812149 | G | A | 28 | 1 | AT1G11330 | S-locus lectin protein kinase family protein | G533S |
| | 3898329 | G | A | 25 | 1 | AT1G11593 | Plant invertase/pectin methylesterase inhibitor superfamily protein | G110R |
| | 4122339 | G | A | 26 | 0.92 | AT1G12140 | flavin-monooxygenase glucosinolate S-oxygenase 5 | intron |
| | 4344329 | G | A | 25 | 0.96 | AT1G12740 | CYP87A2 | D424N |
| | 4568312 | G | A | 34 | 1 | AT1G13330 | HOMOLOGOUS-PAIRING PROTEIN 2 | intron |
| | 4860480 | G | A | 25 | 0.96 | AT1G14220 | Ribonuclease T2 family protein | -878 |
| L-19 (*rda2-7*) | 3019541 | G | A | 34 | 0.97 | AT1G09350 | galactinol synthase 3 | -348 |
| | 3193280 | G | A | 20 | 1 | AT1G09830 | Glycinamide ribonucleotide synthetase | intron |
| | 3214713 | G | A | 26 | 1 | AT1G09890 | Rhamnogalacturonate lyase family protein | intron |
| | 3232906 | G | A | 22 | 1 | AT1G09932 | Phosphoglycerate mutase family protein | L7F |
| | 3284674 | G | A | 23 | 0.95 | AT1G10060 | branched-chain amino acid transaminase 1 | G27E |
| | 3508715 | G | A | 24 | 1 | AT1G10610 | basic helix-loop-helix (bHLH) DNA-binding superfamily protein | R430K |
| | 3536634 | G | A | 24 | 0.91 | AT1G10670 | ATP-citrate lyase A-1 | intron |
| | 3652199 | G | A | 23 | 1 | AT1G10930 | RECQ4A | R455* |
| | 3810696 | G | A | 13 | 0.92 | AT1G11330 | S-locus lectin protein kinase family protein | A109T |
| | 4125184 | G | A | 24 | 0.95 | AT1G12150 | Plant protein of unknown function (DUF827) | P49S |
| | 4241838 | G | A | 21 | 1 | AT1G12440 | A20/AN1-like zinc finger family protein | A162V |

| | | | | | | | | |
|---|---|---|---|---|---|---|---|---|
| L-31 (*rda2-5*) | 2048081 | G | A | 16 | 1 | AT1G06680 | PSBP-1 | D48N |
| | 2379544 | G | A | 32 | 0.96 | AT1G07700 | Thioredoxin superfamily protein | -671 |
| | 3129986 | G | A | 30 | 0.93 | AT1G09660 | RNA-binding KH domain-containing protein | intron |
| | 3812904 | G | A | 25 | 1 | AT1G11330 | S-locus lectin protein kinase family protein | E700K |
| | 3853811 | G | A | 21 | 1 | AT1G11450 | nodulin MtN21-like transporter family protein | G35D |
| | 4424549 | G | A | 20 | 1 | AT1G12970 | plant intracellular ras group-related LRR 3 | intron |
| | 4982756 | G | A | 22 | 0.95 | AT1G14560 | CoA Carrier 1 | intron |
| L-46 (*rda2-8*) | 3171136 | G | A | 37 | 0.94 | AT1G09790 | COBRA-like protein 6 precursor | -317 |
| | 3292819 | G | A | 24 | 1 | AT1G10090 | Early-responsive to dehydration stress protein | P323S |
| | 3402979 | G | A | 31 | 0.96 | AT1G10385 | Vps51/Vps67 family protein | -246 |
| | 3696534 | G | A | 34 | 1 | AT1G11080 | serine carboxypeptidase-like 31 | intron |
| | 3811035 | G | A | 31 | 1 | AT1G11330 | S-locus lectin protein kinase family protein | G222R |
| | 4004066 | G | A | 22 | 1 | AT1G11870 | Seryl-tRNA synthetase | D58N |
| | 4160382 | G | A | 43 | 1 | AT1G12250 | Pentapeptide repeat-containing protein | intron |
| | 4187742 | G | A | 24 | 0.95 | AT1G12310 | Calcium-binding EF-hand family protein | L69F |
| | 4465317 | G | A | 36 | 1 | AT1G13100 | CYP71B29 | D418N |
| | 4543353 | G | A | 28 | 0.96 | AT1G13260 | ETHYLENE RESPONSE DNA BINDING factor 4 | R323K |
| | 4958922 | G | A | 21 | 0.95 | AT1G14490 | Putative AT-hook DNA-binding family protein | P135L |
| L-55 (*rda2-6*) | 2675350 | G | A | 28 | 1 | AT1G08465 | YABBY2 | -684 |
| | 2850878 | G | A | 16 | 1 | AT1G08890 | Major facilitator superfamily protein | junction |
| | 2935539 | G | A | 23 | 1 | AT1G09090 | respiratory burst oxidase homolog B | W606* |
| | 2971223 | G | A | 21 | 1 | AT1G09195 | Ppx-GppA phosphatase | L79F |
| | 3109014 | G | A | 33 | 1 | AT1G09600 | Protein kinase superfamily protein | G133E |
| | 3169313 | G | A | 25 | 1 | AT1G09790 | COBRA-like protein 6 precursor | intron |
| | 3265340 | G | A | 36 | 1 | AT1G10000 | Ribonuclease H-like superfamily protein | -552 |
| | 3710015 | G | A | 19 | 1 | AT1G11110 | LisH and RanBPM domains containing protein | -350 |
| | 3812391 | G | A | 19 | 0.94 | AT1G11330 | S-locus lectin protein kinase family protein | G585R |
| | 3822692 | G | A | 21 | 1 | AT1G11360 | Adenine nucleotide alpha hydrolases-like superfamily protein | L70F |
| L-66 (*rda2-6*) | 1190279 | G | A | 23 | 1 | AT1G04410 | cytosolic-NAD-dependent malate dehydrogenase 1 | T136I |
| | 1506968 | G | A | 27 | 1 | AT1G05200 | glutamate receptor 3.4 | G402R |
| | 1803066 | G | A | 29 | 0.93 | AT1G05940 | cationic amino acid transporter 9 | A238V |
| | 2175774 | G | A | 28 | 0.96 | AT1G07090 | LIGHT SENSITIVE HYPOCOTYLS 6 | -982 |
| | 2263950 | G | A | 35 | 1 | AT1G07370 | PROLIFERATING CELLULAR NUCLEAR ANTIGEN 1 | junction |
| | 2675350 | G | A | 27 | 0.92 | AT1G08465 | YABBY2 | -684 |
| | 2935539 | G | A | 32 | 0.93 | AT1G09090 | respiratory burst oxidase homolog B | W606* |
| | 2971223 | G | A | 28 | 0.92 | AT1G09195 | Ppx-GppA phosphatase | L79F |
| | 3109014 | G | A | 36 | 0.94 | AT1G09600 | Protein kinase superfamily protein | G133E |
| | 3169313 | G | A | 30 | 0.93 | AT1G09790 | COBRA-like protein 6 precursor | intron |
| | 3265340 | G | A | 39 | 0.97 | AT1G10000 | Ribonuclease H-like superfamily protein | -552 |

| | | | | | | | | |
|---|---|---|---|---|---|---|---|---|
| | 3710015 | G | A | 28 | 0.96 | AT1G11110 | LisH and RanBPM domains containing protein | -350 |
| | 3812391 | G | A | 26 | 1 | AT1G11330 | S-locus lectin protein kinase family protein | G585R |
| | 3822692 | G | A | 27 | 1 | AT1G11360 | Adenine nucleotide alpha hydrolases-like superfamily protein | L70F |
| L-74 (*rda2-6*) | 2675350 | G | A | 35 | 0.97 | AT1G08465 | YABBY2 | -684 |
| | 2850878 | G | A | 20 | 1 | AT1G08890 | Major facilitator superfamily protein | junction |
| | 2935539 | G | A | 32 | 1 | AT1G09090 | respiratory burst oxidase homolog B | W606* |
| | 2971223 | G | A | 26 | 0.92 | AT1G09195 | Ppx-GppA phosphatase | L79F |
| | 3109014 | G | A | 23 | 0.95 | AT1G09600 | Protein kinase superfamily protein | G133E |
| | 3169313 | G | A | 32 | 0.96 | AT1G09790 | COBRA-like protein 6 precursor | intron |
| | 3265340 | G | A | 32 | 0.96 | AT1G10000 | Ribonuclease H-like superfamily protein | -552 |
| | 3710015 | G | A | 20 | 1 | AT1G11110 | LisH and RanBPM domains containing protein | -350 |
| | 3812391 | G | A | 38 | 1 | AT1G11330 | S-locus lectin protein kinase family protein | G585R |
| | 3822692 | G | A | 26 | 1 | AT1G11360 | Adenine nucleotide alpha hydrolases-like superfamily protein | L70F |

| Mutant lines | Chromosome coordinate | Reference base | Altered base | Depth | SNP-index | Accession | Note | Type of mutation |
|---|---|---|---|---|---|---|---|---|
| L-53 (*ncer2-2*) | 14610412 | C | T | 15 | 1 | AT2G34660 | multidrug resistance-associated protein 2 | intron |
| | 15000374 | C | T | 23 | 1 | AT2G35690 | acyl-CoA oxidase 5 | intron |
| | 15458693 | C | T | 11 | 1 | AT2G36850 | glucan synthase-like 8 | intron |
| | 15681384 | C | T | 18 | 1 | AT2G37370 | centrosomal protein of 135 kDa-like protein | intron |
| | 15835137 | C | T | 27 | 1 | AT2G37770 | Chloroplastic aldo-keto reductase | intron |
| | 15907123 | C | T | 14 | 1 | AT2G38010 | Neutral/alkaline non-lysosomal ceramidase | Q88* |
| | 15946051 | C | T | 21 | 1 | AT2G38090 | Duplicated homeodomain-like superfamily protein | intron |
| | 16082533 | C | T | 18 | 1 | AT2G38400 | ALANINE:GLYOXYLATE AMINOTRANSFERASE 2 | -1247 |
| | 16151194 | G | A | 10 | 1 | AT2G38620 | cyclin-dependent kinase B1;2 | -1358 |
| | 16411647 | C | T | 11 | 1 | AT2G39300 | CAP-gly domain linker | R382K |
| | 16431177 | C | T | 18 | 1 | AT2G39350 | ATP-binding cassette G1 | G407D |
| | 16796718 | C | T | 15 | 1 | AT2G40220 | ABA INSENSITIVE 4 | G289S |
| | 17201011 | C | T | 13 | 0.92 | AT2G41250 | Haloacid dehalogenase-like hydrolase superfamily protein | G273R |
| | 17295694 | C | T | 27 | 0.96 | AT2G41475 | Embryo-specific protein 3 | intron |
| L-107 (*ncer2-3*) | 15425848 | G | A | 21 | 1 | AT2G36810 | GRAVITROPISM 6 | A688V |
| | 15591280 | G | A | 12 | 1 | AT2G37100 | protamine P1 family protein | R45N |
| | 15906997 | G | A | 14 | 1 | AT2G38010 | Neutral/alkaline non-lysosomal ceramidase | G46S |
| | 15992925 | G | A | 13 | 1 | AT2G38170 | cation exchanger 1 | A85V |
| | 16425133 | G | A | 13 | 1 | AT2G39340 | SAC3A | G142E |
| | 17103110 | G | A | 9 | 1 | AT2G40980 | Protein kinase superfamily protein | G255E |
| | 17127561 | G | A | 9 | 1 | AT2G41060 | RNA-binding (RRM/RBD/RNP motifs) family protein | D142N |

Red highlighting indicates the genes mutated in each mutant. Gray highlighting indicates the intronic mutations (intron) and mutations within 2 kb upstream of the transcription start site (position from the start codon).

**Table S3. List of peptides identified using nano-LC-MS/MS and Mascot.**

| Sample no. | Protein | Peptides | Mascot Score (peptide) | Mascot Score (Total) | Mr (kDa) | Sequence Coverage (%) | Identification |
|---|---|---|---|---|---|---|---|
| 1 | Neutral ceramidase 2 | YGELYTEK | 46 | 100 | 83.156 | 5 | NCER2_ARATH |
| | | GDLLDAGVNR | 38 | | | | |
| | | YDVDKEMTLVK | 18 | | | | |
| | | SPSSYLNNPAAER | 74 | | | | |
| 2 | Neutral ceramidase 2 | MAVELFNK | 40 | 183 | 83.156 | 10 | NCER2_ARATH |
| | | MAVELFNK | 34 | | | | |
| | | SFLISSDPK | 27 | | | | |
| | | GPQPPDLLDK | 36 | | | | |
| | | YEGASTLYGR | 49 | | | | |
| | | VPESAVAGVYR | 55 | | | | |
| | | LATALVNGLTLPR | 83 | | | | |
| | | QISLLSPVVVDSTPLGVK | 26 | | | | |

The proteins identified in each band are shown (a total Mascot Score >50). The identified peptides with their Mascot Score, the calculated molecular mass based on the amino-acid sequence, and the sequence coverage are shown. M = oxidized M.

#### Table S4. Primers used in this study.

| Name | Sequence (5'-3') | Description |
|---|---|---|
| WRKY33_qPCR_F | CTCAAGCACCATATACACTTCA | For qRT-PCR |
| WRKY33_qPCR_R | CCTTTGCTCTAGAGAATCCACC | |
| AtFRK1_qPCR_F | ATCTTCGCTTGGAGCTTCTC | |
| AtFRK1_qPCR_R | TGCAGCGCAAGGACTAGAG | |
| At1g51890_qPCR_F | CCAGTTTGTTCTGTAATACTCAGG | |
| At1g51890_qPCR_R | CTAGCCGACTTTGGGCTATC | |
| UBC_qPCR_F | CTGCGACTCAGGGAATCTTCTAA | |
| UBC_qPCR_R | TTGTGCCATTGAATTGAACCC | |
| pWRKY33_F-SalI | TTTAAACGAATTCGCGTCGACCCTCCGTAGCTTCC TTCTCGTTCTTTG | For pBIB-HYG-pWRKY33-LUC |
| pWRKY33_R | TTTGGCGTCTTCCATACGAAAAATGGAAGTTTGTT TTATA | |
| LUC_F | ATGGAAGACGCCAAAAACATAAAG | |
| LUC_R-KpnI | GGTACCTTACACGGCGATCTTTCCGCCC | |
| RDA2_genome_F | ATGCCTGCAGGTCGAGAATCGTTGTTTCAGAACAT TTGGG | For pBI101-RDA2 (genomic) |
| RDA2_genome_R | GATCGGGGAAATTCGTTAACGTCCTGTTACAGCTG TGAGG | |
| NCER2_genome_F | ATGCCTGCAGGTCGAACTGCCTGTAAGAAATTACA TAACG | For pBI101-NCER2 (genomic) |
| NCER2_genome_R | GATCGGGGAAATTCGTCATACAACTACAAAGGCAC TAGAAG | |
| pNCER2_F | ATGCCTGCAGGTCGAACTGCCTGTAAGAAATTACA TAACG | For pBIB-KAN-HA-NCER2 |
| pNCER2_R | GGTTGATTGAAGTGAGCTTGAGAAGGG | |
| NCER2-SP_F | TCACTTCAATCAACCATGGCTGTTTCACTACCATTG TTTC | |
| SP-HA_R | TAGTCAGGAACATCGTATGGGTATGCATAAACCGTT CTTGAAAGAAG | |
| HA-NCER2_F | CATACGATGTTCCTGACTATGCGTACTTAATCGGAG TCGGAAGCTATG | |
| NCER2_orf_R | GATCGGGGAAATTCGTCATACAACTACAAAGGCAC TAGAAG | |
| pRDA2_F | ATGCCTGCAGGTCGAGAATCGTTGTTTCAGAACAT TTGGG | For pBIB-KAN-RDA2-3xHA |
| pRDA2_R | AACTCTTTGAAACAGAGAATGACTCC | |
| RDA2_orf_F | CGGTACCCGGGGATCCATGGTGGTTTCAGTGACCA TACG | |
| RDA2_orf_R | ACGTCCTGTTACAGCTGTGAGGCTC | |
| 3xHA-F | ACAGCTGTAACAGGACGTGGATCCTACCCATACGA TGT | |
| 3xHA-R | GATCGGGGAAATTCGAGAGTTAAAGGCCTCGAGTT AAGCG | |
| pEAQ-HA-NCER2_F | CAAATTCGCGACCGGTATGGCTGTTTCACTACCATT GTTT | For pEAQ-HA-NCER2 |
| pEAQ-HA-NCER2_stop_R | TACCCCCGGGGTCGACTCATACAACTACAAAGGCA CTAG | |
| NCER2-G46S_bottom_F | CAACATGATGAGCTACGCAAACTC | For pEAQ-HA-NCER2$^{G46S}$ |
| NCER2-G46S_top_R | GCGTAGCTCATCATGTTGACATCAGC | |

#### Table S5. Summary of next-generation sequencing of W33-1B and 11 mutant lines.

| Name | No. of bulked plant | No. of PE reads | Read size (bp) | Total size (Gbp) | Coverage (%) | Read depth | Purpose |
|---|---|---|---|---|---|---|---|
| W33-1B | - | 21,372,837 | 150 | 6.5 | 99.86 | 36.28 | To make consensus genome |
| L-09 | 30 | 23,984,219 | 150 | 7.2 | 99.86 | 42.70 | To perform MutMap analysis |
| L-12 | 30 | 20,851,955 | 150 | 6.3 | 99.86 | 37.92 | To perform MutMap analysis |
| L-16 | 30 | 20,928,591 | 150 | 6.3 | 99.86 | 37.36 | To perform MutMap analysis |
| L-19 | 30 | 20,573,214 | 150 | 6.2 | 99.86 | 36.60 | To perform MutMap analysis |
| L-31 | 30 | 19,203,463 | 150 | 5.8 | 99.86 | 33.80 | To perform MutMap analysis |
| L-46 | 30 | 22,464,100 | 150 | 6.8 | 99.86 | 40.60 | To perform MutMap analysis |
| L-55 | 30 | 20,906,642 | 150 | 6.3 | 99.86 | 35.34 | To perform MutMap analysis |
| L-66 | 30 | 21,848,777 | 150 | 6.6 | 99.86 | 40.86 | To perform MutMap analysis |
| L-74 | 30 | 23,108,408 | 150 | 7.0 | 99.86 | 42.06 | To perform MutMap analysis |
| L-53 | 30 | 16,062,165 | 150 | 4.8 | 99.86 | 26.96 | To perform MutMap analysis |
| L-107 | 30 | 16,538,531 | 150 | 5.0 | 99.86 | 26.54 | To perform MutMap analysis |

**Table S6. Number of reliable SNPs detected between wild type W33-1B and each of the 11 mutant lines.**

| Mutant lines | Reference sequence | Number of SNPs detected | | |
|---|---|---|---|---|
| | | Total SNPs and indel | P-value < 0.05 | P-value < 0.01 |
| L-09 | W33-1B | 747 | 87 | 71 |
| L-12 | W33-1B | 944 | 155 | 113 |
| L-16 | W33-1B | 919 | 149 | 88 |
| L-19 | W33-1B | 958 | 189 | 122 |
| L-31 | W33-1B | 479 | 70 | 48 |
| L-46 | W33-1B | 1,113 | 187 | 106 |
| L-55 | W33-1B | 516 | 94 | 71 |
| L-66 | W33-1B | 580 | 63 | 54 |
| L-74 | W33-1B | 587 | 78 | 65 |
| L-53 | W33-1B | 608 | 108 | 71 |
| L-107 | W33-1B | 1,007 | 238 | 152 |
